# Aneuploidy alleviates the cell proliferation defect caused by mutations affecting origin licensing in *Saccharomyces cerevisiae*

**DOI:** 10.1101/2025.04.29.651238

**Authors:** Christophe de La Roche Saint-André

## Abstract

Although aneuploidy is generally detrimental to the survival and growth of normal cells, it can be beneficial under certain stress conditions, such as those caused by harmful mutations. In *Saccharomyces cerevisiae*, we find that the spontaneous doubling of chromosome III in the *orc5-1 set1Δ* mutant accelerates proliferation, a benefit resulting from a reduction in the negative effect of the *orc5-1* mutation. Enhanced proliferation is also observed when a fragment from a different chromosome is introduced, demonstrating that the benefit is not simply due to extra copies of specific genes. A comparable growth-enhancing effect of an extra chromosome is observed for mutations affecting other proteins involved in DNA replication licensing. The suppression of *orc5-1* growth defect is also observed in the absence of the G1 cyclin Cln3, which lengthens the G1 phase, while overexpressing *CLN3*, which shortens G1, has the opposite effect. Additionally, Cln3 loss mirrors the effect of an extra chromosome for other mutations. These findings indicate that the severity of mutations impacting origin licensing hinges on the length of the G1 phase. Thus, we propose that the fitness-enhancing effect of an extra chromosome in DNA replication licensing mutants largely stems from its ability to extend G1, compensating for inefficient origin licensing.

## Introduction

Aneuploidy is a genomic condition characterized by an abnormal number of chromosomes, resulting from errors in chromosome segregation. Such a break in the usual number of chromosomes leads to changes in the number of copies of many genes, which generally has repercussions on the amount of proteins they encode. The result is stoichiometric imbalances in certain protein complexes, leading to an increase in protein misfolding, which in turn causes proteotoxic stress due to an increased demand for protein quality control mechanisms [1–4]. This overall negative effect, which is not specific to the chromosomes whose number is altered, compromises the metabolic activity and genome stability of the cell, and affects both its survival and growth in a favorable environment [5,6]. Of all the negative consequences of aneuploidy, many appear to be shared by all eukaryotic cells. For example, the gain of an extra copy of a chromosome causes a delay in the G1 phase in *Saccharomyces cerevisiae* and a lengthening of the G1 and S phases in mammalian cells, cell cycle modifications that slow down cell proliferation [1,5,7].

The cost of aneuploidy can be offset by certain advantages that enable cells to survive and divide more efficiently when conditions become unfavorable [8,9]. In humans, aneuploidy is considered as a hallmark of cancer cells. It is widely prevalent in most solid tumors and likely contributes to cancer progression [10]. In *Saccharomyces cerevisiae*, a gain of chromosomes can prove beneficial when cells are confronted with different types of stress, whether external, i.e. adverse environments, or internal, such as the consequences of deleterious mutations [11–15]. In such stressful conditions, aneuploidy can enable rapid adaptive evolution at least transiently [16,17]. In numerous cases, the beneficial effect linked to the presence of extra chromosomes results from the elevated expression of specific genes which alleviates the negative effects of the stress, whether it is environmental or genetic [12–16]. For example, chromosomal gain can confer drug resistance in pathogenic fungi by increasing the copy number of genes encoding the drug target and drug efflux transporters [18]. Specific aneuploidies also lead to drug resistance in human cancer cells [19]. Although the drug resistance phenotype may be linked to altered expression of certain dose-sensitive genes, a non-specific effect of aneuploidy may also be involved. As such, the G1 extension associated with aneuploidy could delay the stage of the cell cycle where the drug exerts its antiproliferative effects [20].

We recently reported a role for Set1 H3K4 methylase during DNA replication when S-phase dynamics change in response to decreased origin firing in *Saccharomyces cerevisiae* [21]. In this context, we noticed the spontaneous establishment of a chromosome III disomy when *set1Δ* was combined with the temperature-sensitive allele *orc5-1*, which affects a subunit of the origin replication complex (ORC). This led us to investigate why the chromosome III disomy was selected in this mutant context. Here we show that the presence of an extra chromosome III is associated with better proliferation of the double mutant. Strikingly, this positive effect was reproduced in the presence of fragments of either chromosome III or VII, revealing that it was not specific to any particular chromosome. Our data further showed that the beneficial effects of an extra chromosome are independent of Set1 and present in the *orc5-1* mutant alone, as well as other mutants affecting, directly or indirectly, origin licensing, i.e. the formation of pre-replicative complexes (pre-RCs) at potential origins of replication. Extending the G1 phase either by growing the cells in a less nutrient-rich environment or by inactivating the G1 cyclin Cln3, similarly improves the growth of mutants affecting the pre-RC complex. Taken together, we propose that the positive effect of an extra chromosome is likely caused by the lengthening of the G1 phase, which compensates for the defect in origin licensing associated with different mutations.

## Results

### Chromosome III disomy stimulates cell proliferation in the *orc5-1 set1Δ* mutant

In the course of a previous study [21], we constructed a diploid strain in the SK1 genetic background, homozygous for the *orc5-1* and *set1Δ* mutations and with both chromosomes III carrying a different antibiotic resistance marker (*Hph*MX or *Nat*MX) near their centromeres (S1A Fig). Strikingly, after sporulation of the diploid, an excess of one marker (*Nat*MX) was found in a large fraction of tetrads, with half of the spores carrying both markers (S1A Fig). This suggested that two copies of chromosome III were in these spores, which was confirmed by pulsed-field gel electrophoresis (PFGE) analysis (S1B Fig). No duplication was observed in tetrads from diploids carrying no or single *orc5-1* or *set1Δ* mutation (not shown). We concluded that cells disomic for chromosome III arose spontaneously and were selected for in one of the *orc5-1 set1Δ* haploids that was used to generate the diploid.

We noticed that *orc5-1 set1Δ* spore colonies with two chromosomes III were larger in size (S1A Fig). This was confirmed by crossing a disomic *orc5-1 set1Δ* haploid with a wild-type, to generate a heterozygous *ORC5/orc5-1 SET1/set1Δ* diploid with three chromosomes III (Fig 1A, left). After sporulation, spores with all four genotypes (WT, *set1Δ*, *orc5-1*, *orc5-1 set1Δ*) carrying one or two chromosomes III were isolated on YPD plates and incubated at 25°C for 3 days, before the area of the spore-derived colonies was measured (Fig 1A, right). In the presence of a single chromosome III, colonies corresponding to *orc5-1 set1Δ* spores were the smallest, in line with published data [21]. Chromosome III disomy was associated with a significant reduction in the size of colonies derived from wild-type spores, which was consistent with the proliferation defect described in the presence of an extra chromosome [1]. In contrast, an extra chromosome III was associated with significantly larger *orc5-1 set1Δ* colonies. No impact on colony size was observed in single mutants at 25°C, a permissive temperature for *orc5-1*.

**Fig 1.**
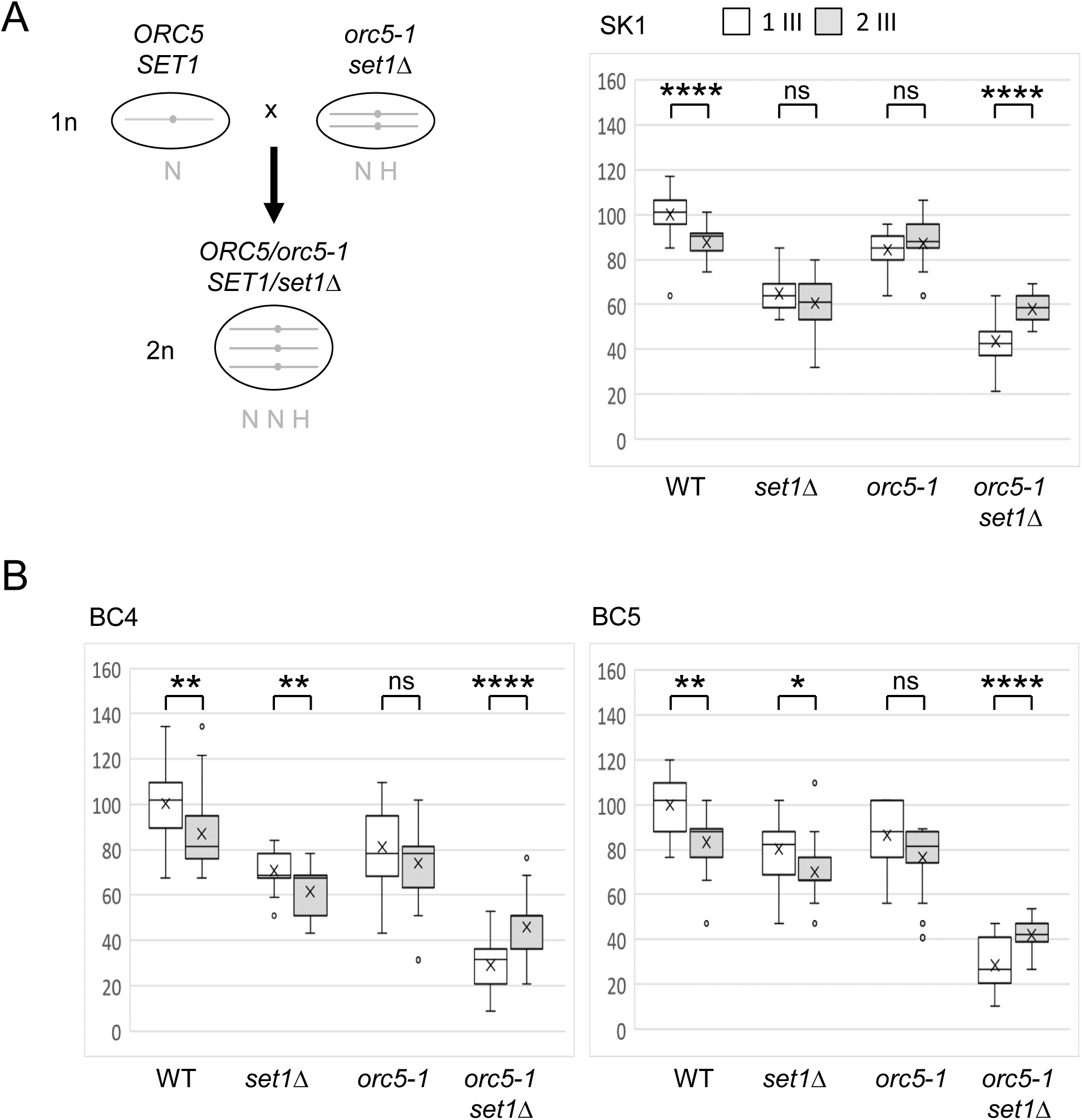
A growth advantage is associated with chromosome III disomy in *orc5-1 set1Δ*. (A) Left : a *ORC5/orc5-1 SET1/set1Δ* diploid (2n) with three III chromosomes (SK1 background) was obtained by crossing a WT haploid (1n), bearing one chromosome III with the *Nat*MX centromeric marker (N), with an *orc5-1 set1Δ* haploid (1n), bearing two chromosomes III with the *Nat*MX or *Hph*MX centromeric markers (NH). Right : box plots corresponding to sizes of the spore-derived colonies grown 3 days at 25° on YPD are shown according to the spore genotype and the number, one or two, of chromosome III (white and grey boxes respectively). Sizes were normalized by setting the mean size of the WT with one chromosome III at 100. P-values in a one-way ANOVA test are indicated : ns *p* > 0.05, * *p* < 0.05, ** *p* < 0.01, *** *p* < 0.001, **** *p* < 0.0001. (B) Same box plot presentations corresponding to the fourth (BC4, left) and fifth (BC5, right) backcrosses from the SK1 background to the W303 background.

A larger colony can result from either larger cells, due to a delayed cell division, or from more cells, due to faster cell proliferation. To determine which of these alternatives explains the effect of chromosome III disomy on *orc5-1 set1Δ* colony size, we performed a competitive growth experiment (S2 Fig). *orc5-1 set1Δ* cells having one or two chromosomes III with one centromeric marker (*Nat*MX) were cocultured at a 1/1 ratio with cells having a single chromosome III with *Hph*MX. The evolution of the *Nat*MX/*Hph*MX cell ratio was followed for several days in liquid culture at 25°C. While the ratio was maintained in the control as expected, it increased in favor of cells with two chromosomes III, with a ratio of around 100/1 after twenty cell population doublings. This proliferation advantage linked to the presence of two chromosomes III was therefore responsible for the larger size of the *orc5-1 set1Δ* colonies.

To check that the positive effect of chromosome III disomy in *orc5-1 set1Δ* was not a particular feature of the SK1 genetic background, we performed backcrosses to introduce the two genetically marked chromosomes III into the W303 genetic background. We assessed by PFGE analysis that the extra copy of chromosome III was stably transmitted during the backcrosses (S3 Fig). The effect of chromosome III disomy on spore colony growth was tested at the fourth and fifth backcrosses (Fig 1B). In both cases, chromosome III disomy was associated with a growth advantage specifically in *orc5-1 set1Δ*. Thus, the proliferation advantage conferred by chromosome III disomy in *orc5-1 set1Δ* was not restricted to a particular genetic background.

### The positive effect of chromosome III disomy on *orc5-1 set1Δ* cannot be attributed to specific genes

In order to determine whether a specific part of chromosome III was responsible for its positive effect on *orc5-1 set1Δ*, we tested the impact of a fragment (CFIII) corresponding to the left arm of the chromosome [24]. On the right side of the CFIII centromere is an auxotrophic marker (*URA3*) and a mutant tRNA (*SUP11*) able to suppress the ochre stop codon present in the *ade2-1* allele (Fig 2). Cells with *ade2-1* are auxotroph for adenine and accumulate a red pigment when adenine is limiting. On YPD plates not supplemented with adenine, the presence of CFIII is indicated by the white color of the colonies, due to the suppression of the *ade2-1* mutation, whereas on adenine-supplemented YPD medium, the presence of the *URA3* marker is used (Fig 2). The effect of CFIII on colony size was similar to that of the whole chromosome III, whether adenine was limiting or not, with a positive effect on spore colony growth only in the case of *orc5-1 set1Δ*. Therefore, the right arm of chromosome III appeared dispensable for its positive effect on *orc5-1 set1Δ* fitness.

**Fig 2.**
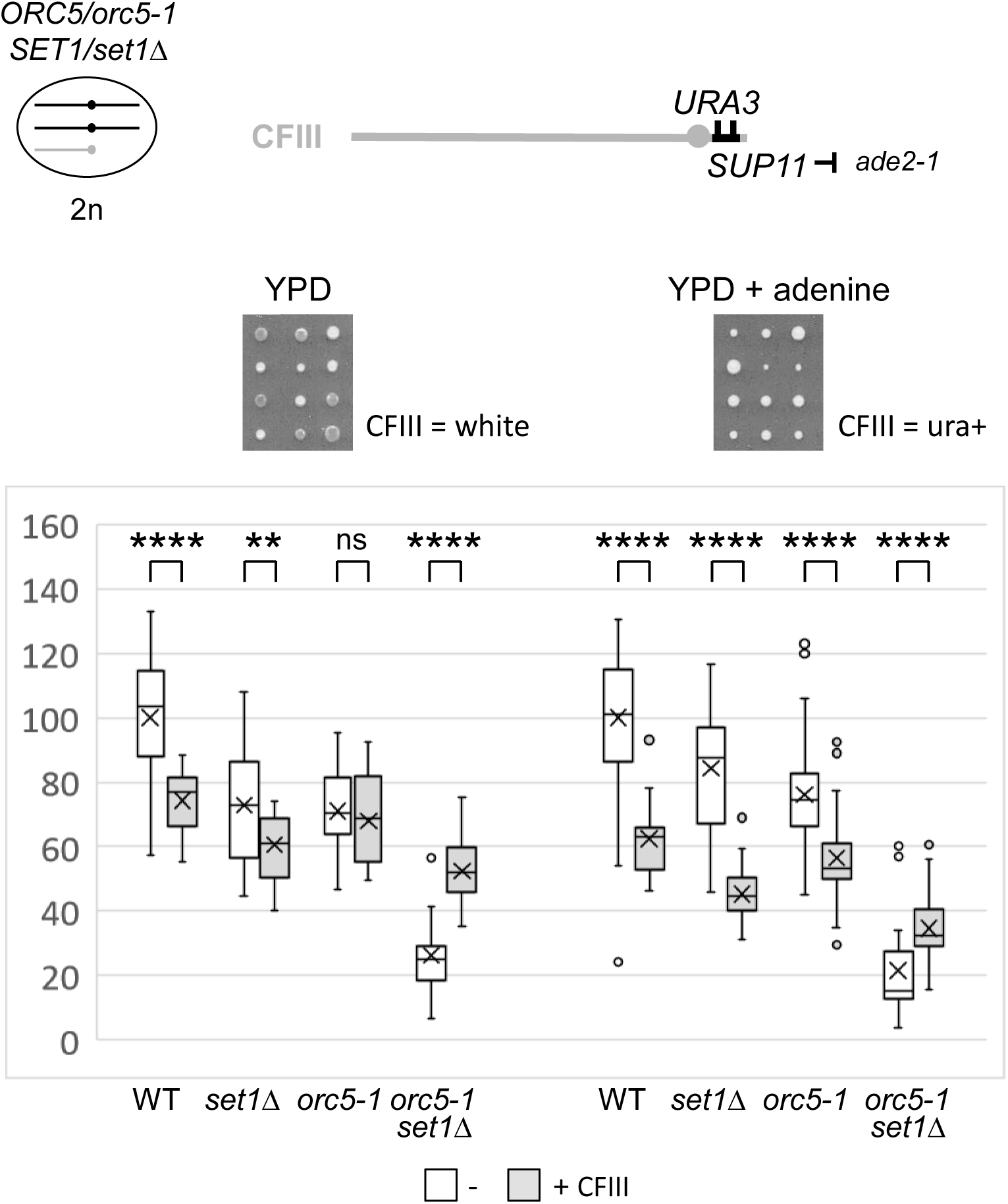
A growth advantage is associated with an extra left arm of chromosome III in *orc5-1* set1Δ. Top left : the *ORC5/orc5-1 SET1/set1Δ* diploid (2n) with two chromosomes III (in black) and the additional chromosome III fragment (CFIII in grey). Top right : schematic drawing of CFIII with its *URA3* and *SUP11* genetic markers (in black). Middle : representative tetrads dissected from the sporulated diploid on standard YPD plates (left) or YPD plates supplemented with adenine (right), grown 3 days at 25°. Presence of CFIII was checked by the white colony color and/or the uracil prototrophy. Bottom: the box plots correspond to sizes of the spore-derived colonies according to the spore genotype and the absence/presence of CFIII. sizes were normalized by setting the mean size of the WT without CFIII at 100. P-values in a one-way ANOVA test are indicated : ns *p* > 0.05, * *p* < 0.05, ** *p* < 0.01, *** *p* < 0.001, **** *p* < 0.0001.

The positive effect associated with the presence of an extra chromosome III or CFIII in *orc5-1 set1Δ* could result from the overexpression of one or more proteins encoded by the left arm of chromosome III, which could exert their positive effect by mitigating the replicative stress present in *orc5-1 set1Δ* [21]. Among the 71 ORFs located on the left arm of chromosome III, two genes were noticed, *MRC1* and *DCC1*. Mrc1 is an S-phase checkpoint protein that stabilizes stalled replication forks, while the Dcc1 protein is required for sister chromatid cohesion establishment behind replication forks [25]. These two genes are involved in genetic interactions both with Set1 complex and ORC genes [21,25,26]. It was therefore conceivable that overexpression of Mrc1 or Dcc1 could be responsible for the positive effect of chromosome III left arm duplication in *orc5-1 set1Δ*.

We thus tested whether inactivating one copy of the *MRC1* or the *DCC1* genes reduced the fitness increase associated with chromosome III disomy in *orc5-1 set1Δ*. The *mrc1Δ* and *dcc1Δ* mutations were introduced through additional backcrosses into the W303 background. In the presence of a single chromosome III, *orc5-1* and *mrc1Δ* were synthetically lethal, as previously described [26] (S4A Fig). In the presence of a second chromosome III, the lethality was suppressed, as expected, and neither the negative nor the positive effect associated with chromosome III disomy on WT or *orc5-1 set1Δ* respectively was changed. This also held true for the *DCC1* deletion (S4B Fig), with near-synthetic lethality of *orc5-1* and *dcc1Δ* when only one chromosome III was present [26] and no significant impact of *dcc1Δ* in the presence of two chromosomes III, excepted in the case of *set1Δ*. It therefore appears that the *MRC1* and *DCC1* genes do not contribute to the positive effect of chromosome III disomy on *orc5-1 set1Δ*.

### A positive effect that is not chromosome-specific

To further determine whether the positive effect on *orc5-1 set1Δ* was specific to chromosome III, we tested an other chromosomal fragment (CFVII) corresponding to the terminal part of the right arm of chromosome VII [24] (S5 Fig). Surprisingly, the effect of CFVII was similar to that of CFIII, with again a clear positive impact on spore colony growth only in the case of *orc5-1 set1Δ*. Therefore, the positive effect on *orc5-1 set1Δ* fitness was not specific to chromosome III.

The effect of an extra chromosome XII, much larger than chromosome III, was also tested (S6 Fig). In line with the fact that the effect of a disomy correlates roughly with the size of the chromosome [1,27], chromosome XII disomy was associated in WT cells with a much greater negative effect than chromosome III disomy. A greater negative effect was also observed in *set1Δ* and *orc5-1*. The growth of *orc5-1 set1Δ* colonies was significantly slowed by the presence of an extra chromosome XII, albeit to a relatively lesser extent compared with the other genotypes. As the number of ORFs on chromosome XII is three times higher than on chromosome III, we propose that in *orc5-1 set1Δ* the non-specific positive effect of disomy was outweighed by the negative consequences of gene overexpression when the number of genes exceeds a certain threshold.

### The improved cell proliferation is linked to the *orc5-1* mutation and depends on the yeast nature of the extra chromosome

We observed that, despite some experimental variability, the impact of an extra chromosome on *orc5-1* colony size was without significant effect at permissive temperature (see Fig 1,2,4). However, at the semi-permissive temperature of 30°C, we observed a significant positive effect of an extra chromosome in the *orc5-1* single mutant (Fig 3A). This was also the case at 32°C, in an experiment with *rad52Δ* instead of *set1Δ* because *orc5-1 set1Δ* spores did not give colonies at this temperature (Fig 3B). It therefore appeared that the positive effect of CFVII was specific to the *orc5-1* mutation, independently of Set1, and was only apparent above a certain temperature, i.e. when the function of the Orc5-1 protein was sufficiently altered.

**Fig 3.**
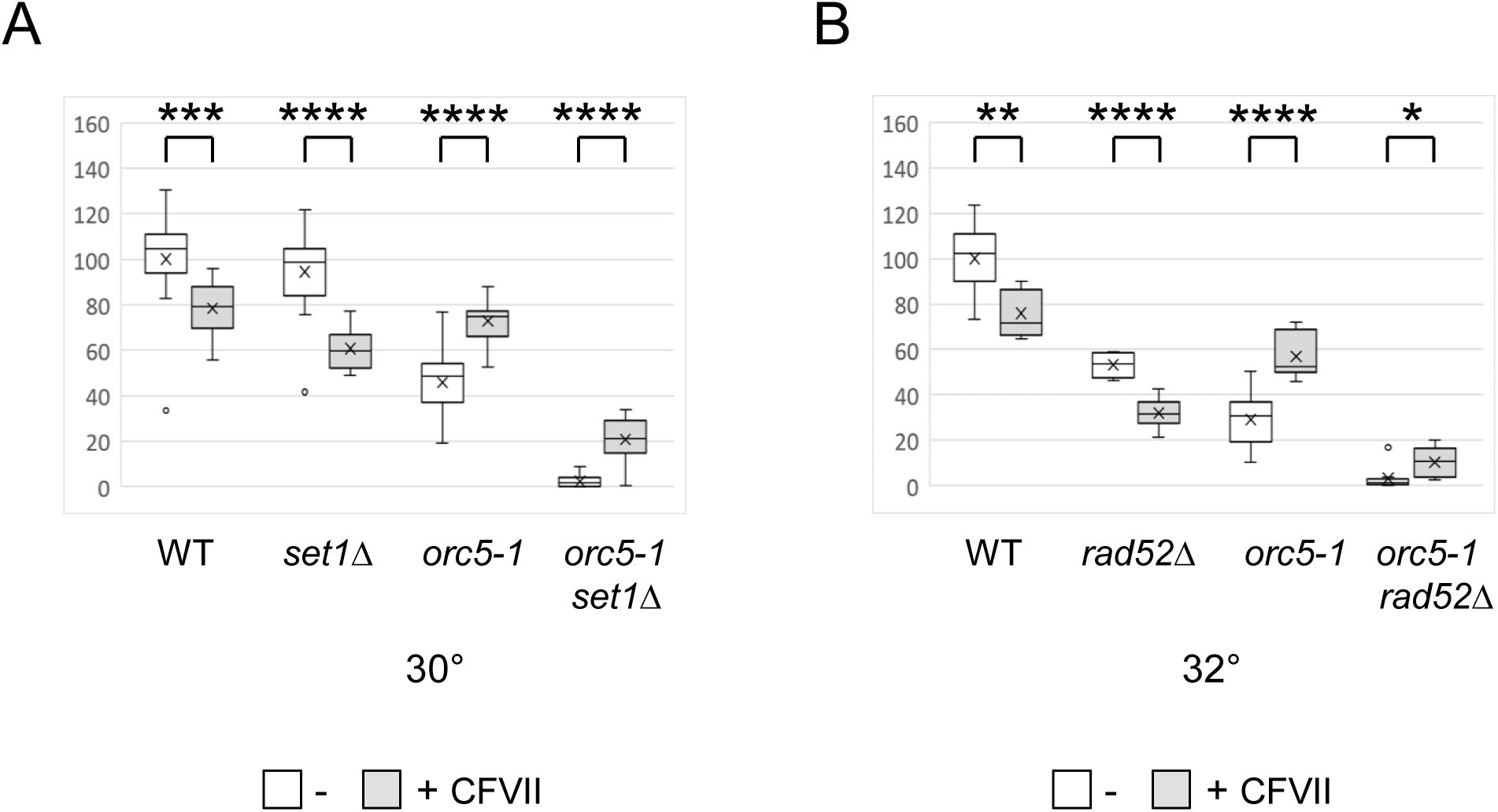
Raising the incubation temperature reveals the positive effect of CFVII on *orc5-1*. (A) Analysis of the meiotic products of the *ORC5/orc5-1 SET1/set1Δ* diploid with CFVII. The box plots correspond to sizes of the spore-derived colonies grown 3 days at 30° on YPD, according to the spore genotype and the absence/presence of CFVII. Sizes were normalized by setting the mean size of the WT without CFVII at 100. P-values in a one-way ANOVA test are indicated : ns *p* > 0.05, * *p* < 0.05, ** *p* < 0.01, *** *p* < 0.001, **** *p* < 0.0001. (B) Same analysis as in (A) with the meiotic products of the *ORC5/orc5-1 RAD52/rad52Δ* diploid with CFVII, excepted that the spore-derived colonies were grown 3 days at 32°.

To ensure that the larger size of *orc5-1* colonies in the presence of an extra chromosome was the result of increased cell proliferation, cells were harvested directly from colonies growing at 32°C and analyzed by flow cytometry. Cell size was determined by measuring forward scatter (FSC). As shown in Figure 4A, in the absence of CFIII, the size of *orc5-1* cells was much larger than that of wild-type cells, suggesting a slowdown in cell proliferation. In line with this, DNA content analysis showed that the proportion of *orc5-1* cells in S and G2/M phases was increased as compared with wild-type cells (Fig 4B). While the presence of CFIII did not influence WT cell size, it was associated with a significant reduction in *orc5-1* cell size (Fig 4A), correlating with a wild-type cell cycle profile (Fig 4B). The fact that *orc5-1* colonies with CFIII contained more cells progressing better through the cell cycle than colonies without CFIII clearly demonstrated improved cell proliferation in the presence of an extra chromosome.

**Fig 4.**
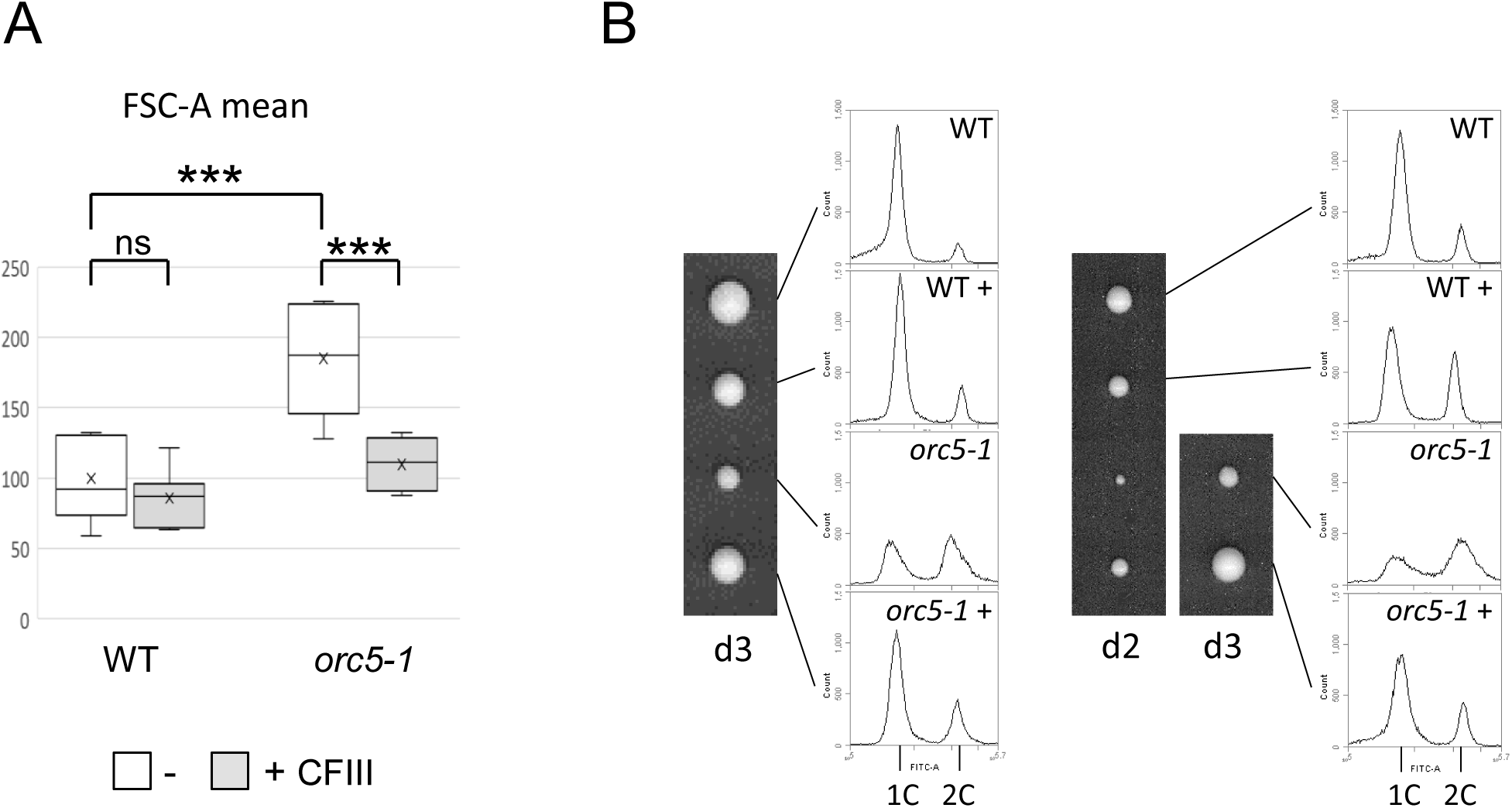
Effect of CFIII on the cell size and the cell cycle. The meiotic products of the *ORC5/orc5-1* diploid with CFIII were grown on YPD at 32°. Cells from spore-derived colonies were analyzed using flow cytometry and their size (A) and DNA content (B) was determined. (A) Box plots corresponding to the average size (FSC-A mean) of the cells according to their genotypes and the absence/presence of CFIII after 3 days of growth. Sizes were normalized by setting the mean size of the WT without CFVII at 100. P-values in a one-way ANOVA test are indicated : ns *p* > 0.05, * *p* < 0.05, ** *p* < 0.01, *** *p* < 0.001, **** *p* < 0.0001. (B) Two tetrads are shown together with DNA content profiles for each spore colony after 2 (d2) or 3 (d3) days of growth. The genotype is indicated as well as the presence of CFIII (+). 1C and 2C indicate the DNA content of cells with unreplicated or fully replicated DNA, respectively.

To determine whether this beneficial effect of an extra chromosome depended on its yeast nature, we examined the effect of human DNA carried by yeast artificial chromosomes (YACs). Introduction of YACs that are 670 kb long (YAC-3) or 850 kb long (YAC-2) had no effect on the wild type (S7 Fig), confirming that YAC strains do not exhibit the proliferation defect commonly associated with an extra chromosome [1,28]. YAC-3 did not suppress the proliferation defect of *orc5-1* and YAC-2 had an additional negative impact. Thus, YACs did not have a positive effect on *orc5-1* growth at semi-permissive temperature (30°C), in contrast with the highly significant effect of CFVII at this temperature. We concluded that this positive effect required a functional yeast chromosome.

The magnitude of the positive effects of CFIII and CFVII on *orc5-1 set1Δ* appeared similar (Fig 2 and S5 Fig). To test if this was also the case for *orc5-1*, and overcome some experimental variability, we compared the effect of the two chromosome fragments in the same experiment. For this purpose, CFIII WT cells were crossed with CFVII *orc5-1* cells and the meiotic products of the resulting diploid were analyzed (S8 Fig). Because the same genetic markers are present on CFIII and CFVII, spores containing either chromosome fragment were genetically indistinguishable, but could be differentiated from spores containing both chromosome fragments (see Materials and methods). As shown in Figure S8, the colony size of spores containing only one chromosomal fragment was homogeneous, indicating that the positive effect of CFIII and CFVII were comparable. In the wild type, as expected, the negative effects of CFIII and CFVII were additive. In *orc5-1*, while either chromosomal fragment exerted a positive effect, a negative effect was associated with the presence of both fragments. This negative effect was nevertheless more limited as compared to the wild type, as shown by the absence of any difference in the size of WT and *orc5-1* colonies in the presence of the two chromosome fragments. These results confirm that in *orc5-1*, a positive effect of an additional chromosome can be detected as long as the amount of added genetic material does not exceed a certain threshold.

### The positive effect of an extra chromosome is not restricted to the *orc5-1* allele

Orc5 is a subunit of the origin recognition complex (ORC) which main role is to direct DNA replication by binding to replication origins. To determine if the positive effect of an extra chromosome was specific to the *orc5-1* mutation, we monitored the impact of CFVII on temperature-sensitive mutants affecting other proteins involved in DNA replication. The incubation temperatures of the dissection plates were adapted to the level of thermosensitivity associated with each mutation.

We first tested a mutation, *orc2-1*, that affects another subunit of the ORC complex (Fig 5A). At 30°C, the growth of *orc2-1* colonies was strongly affected but improved in the presence of CFVII, as already observed for *orc5-1*. The Cdc6 protein interacts with the ORC complex and recruits the MCM complex to replication origins, leading to the formation of pre-replicative complexes (pre-RCs) during G1 phase, a process known as origin licensing [29]. While the presence of CFVII affected wild-type growth, the presence of CFVII had no effect on *cdc6-4* colony size at 36°C (Fig 5B), suggesting that the negative impact of the extra chromosome was offset by a positive effect.

**Fig 5.**
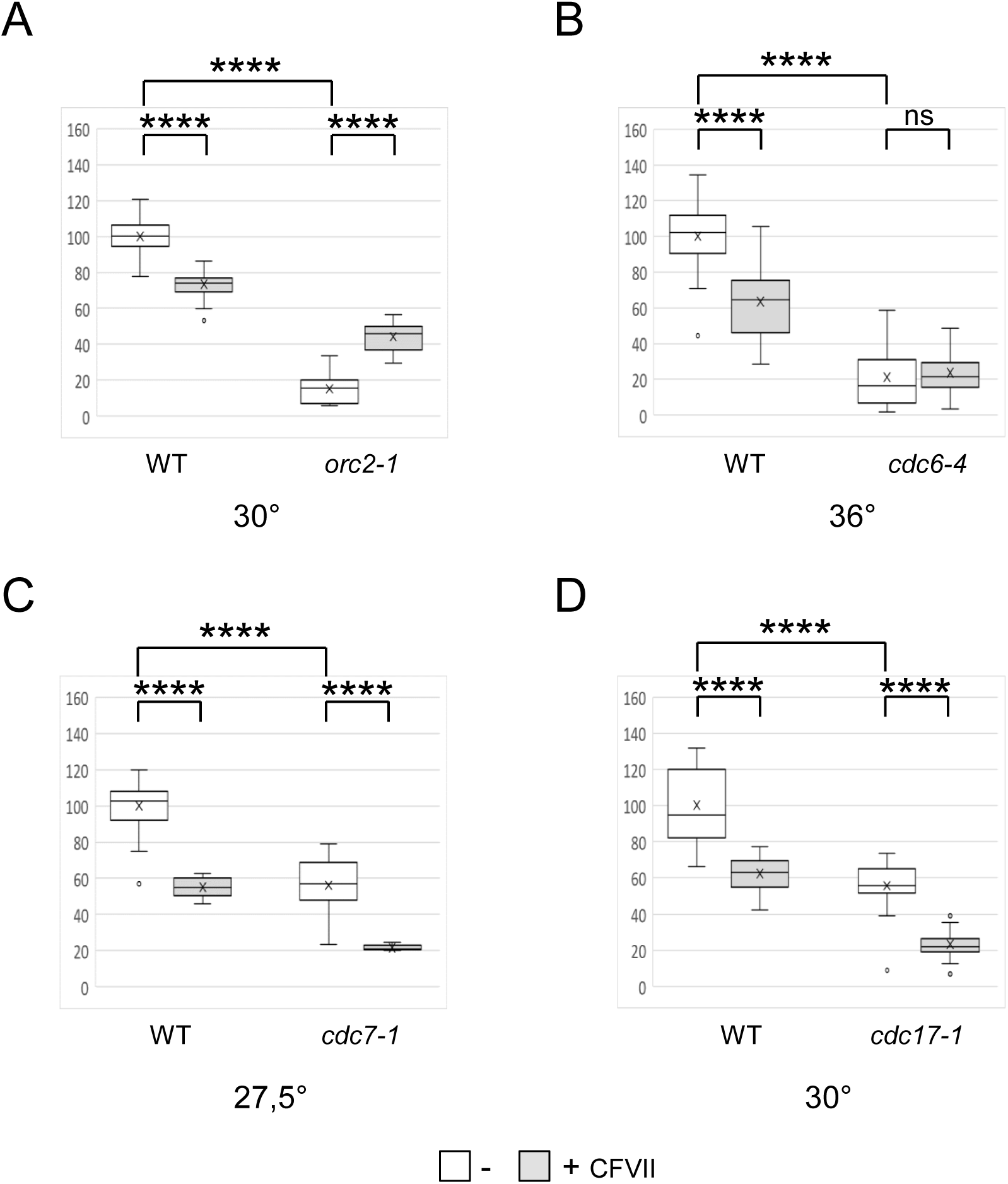
Effect of CFVII on *orc2-1*, *cdc6-4*, *cdc7-1* and *cdc17-1* spore colony size. Analysis of the meiotic products of the *ORC2/orc2-1* (A), *CDC6/cdc6-4* (B), *CDC7/cdc7-1* (C) and *CDC17/cdc17-1* (D) diploids with CFVII. The box plots correspond to sizes of the spore-derived colonies grown 2-3 days at the indicated temperature according to the spore genotype and the absence/presence of CFVII. Sizes were normalized by setting the mean size of the WT without CFVII at 100. P-values in a one-way ANOVA test are indicated : ns *p* > 0.05, * *p* < 0.05, ** *p* < 0.01, *** *p* < 0.001, **** *p* < 0.0001.

We next tested mutations affecting proteins involved in stages subsequent to pre-RC formation. Cdc7, the catalytic subunit of DDK (Dbf4-dependent kinase), is required for origin firing and replication initiation. Cdc17, the catalytic subunit of the DNA polymerase I alpha-primase complex, is essential for the initiation and progression of DNA replication. CFVII was associated with a strong negative effect on both *cdc7-1* (Fig 5C) and *cdc17-1* (Fig 5D) colony growth.

Sister chromatid cohesion (SCC) establishment is coupled to DNA replication and follows the loading of the cohesin complex onto chromatin during the G1 phase. Strong synthetic interactions were demonstrated between *orc5-1* or *orc2-1* and mutations in genes that contribute to SCC [26]. We tested the effect of CFVII on mutations affecting proteins belonging to the cohesin complex (Smc1 and Smc3) or the cohesin loader (Scc2). As expected for mutations affecting SCC, CFVII was unstable and rapidly lost in colonies derived from *smc3-42*, *smc1-259* and *scc2-4* temperature-sensitive spores. Nevertheless, the size of these mixed colonies was still significantly smaller than that of colonies derived from mutant spores without CFVII (S9 Fig). Thus, the presence of CFVII was associated with a negative effect in an SSC mutant context, the magnitude of which was underestimated due to the frequent loss of the chromosomal fragment.

Finally, we evaluated the effect of CFVII on mutations affecting proteins involved in the late stage of the cell cycle, namely the Cdc14 phosphatase and the Cdc15 kinase which are essential for mitotic exit. CFVII was associated with a positive effect on colony growth of both *cdc14-3* and *cdc15-2* (Fig 6). Sic1, an inhibitor of Cdk activity, is one of the substrates that is dephosphorylated by Cdc14 to promote exit from mitosis. The growth of *sic1Δ* colonies was strongly affected by the presence of CFVII (Fig 6), showing that not all mutations of proteins involved in mitotic exit were affected in the same way.

**Fig 6.**
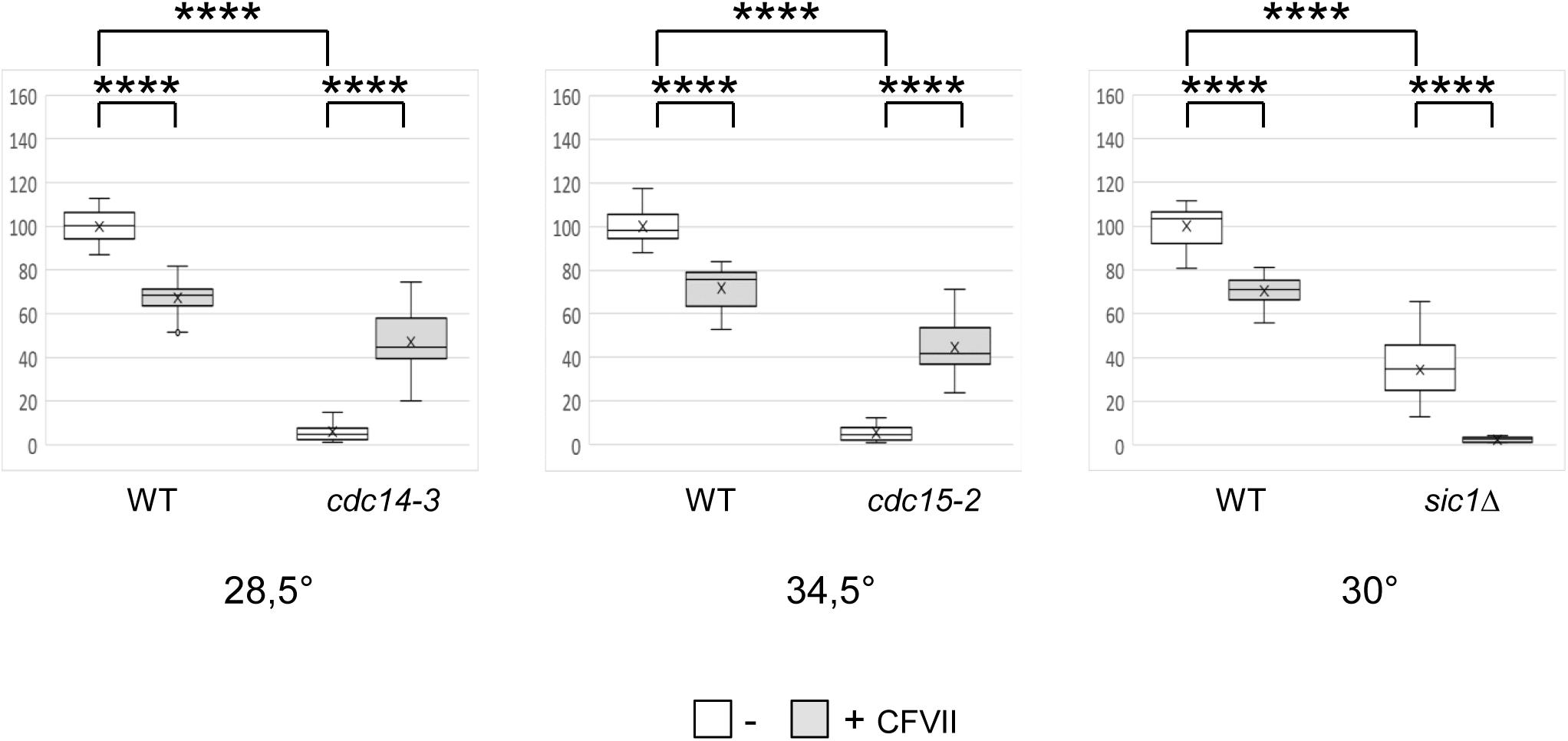
Effect of CFVII on *cdc14-3*, *cdc15-2* and *sic1Δ* spore colony size. Analysis of the meiotic products of the *CDC14/cdc14-3* (left), *CDC15/cdc15-2* (middle) and *SIC1/sic1Δ* (right) diploids with CFVII. The box plots correspond to sizes of the spore-derived colonies grown 2-3 days at the indicated temperature according to the spore genotype and the absence/presence of CFVII. Sizes were normalized by setting the mean size of the WT without CFVII at 100. P-values in a one-way ANOVA test are indicated : ns *p* > 0.05, * *p* < 0.05, ** *p* < 0.01, *** *p* < 0.001, **** *p* < 0.0001.

### A longer G1 phase explains the positive effect of an extra chromosome

As the lengthening of the G1 phase is one of the general consequences of the presence of an extra chromosome [1,27] we hypothesized that it might be responsible for the improved fitness observed in some mutants. In this hypothesis, we would expect to observe a similar effect by prolonging G1 by other means.

Differences in G1 duration explain most of the differences in total cell cycle length, or generation time, between the same cells growing in different media [30]. To increase the duration of G1, spores were thus isolated on the synthetic defined (SD) medium, which is nutritionally poorer than YPD. The deleterious effect of an extra chromosome (CFIII or CFVII) disappeared in WT cells grown on SD (S10A Fig). The negative effect in presence of both CFIII and CFVII, although significant, was relatively weaker than on YPD. It therefore appears that lengthening the G1 phase in SD minimizes the additional effect of an extra chromosome. The effect of the SD medium was then tested on *orc5-1* (S10B Fig). At 30°C, the negative impact of the *orc5-1* mutation on colony growth was attenuated on SD compared to YPD, and the positive effect of CFVII on *orc5-1* colony growth was no longer visible. The incubation temperature had to be increased to 33.5°C, and the growth rate accelerated, for a positive effect of CFVII to be observed again. These results were consistent with the idea that the extra chromosome exerts its positive effect on *orc5-1* via a lengthening of the G1 phase.

We next modified the duration of G1 by deleting the *CLN3* gene. The Cln3 cyclin accumulates during G1 and activates the Cdc28 kinase to promote the G1 to S phase transition [31]. It has been shown that the extension of G1 phase induced by an extra chromosome is associated with a delay in Cln3 accumulation [1,27]. Remarkably, the impact of Cln3 loss was similar to that of an extra chromosome, with a negative effect on wild-type colony size and a positive effect on *orc5-1* colony size (Fig 7A). We investigated whether *CLN3* overexpression had an opposite effect using the *GAL1-CLN3* construct which allows overproduction of Cln3 in the presence of galactose [32]. As expected, the positive effect of CFVII on *orc5-1* was not impacted by galactose (S11A Fig) and the growth defect due to *cln3Δ* was complemented by the *GAL1-CLN3* construct (S11B Fig). Notably, decreasing the length of the G1 phase through *CLN3* overexpression had a negative effect on the growth of *orc5-1* colonies (S11C Fig).

**Fig 7.**
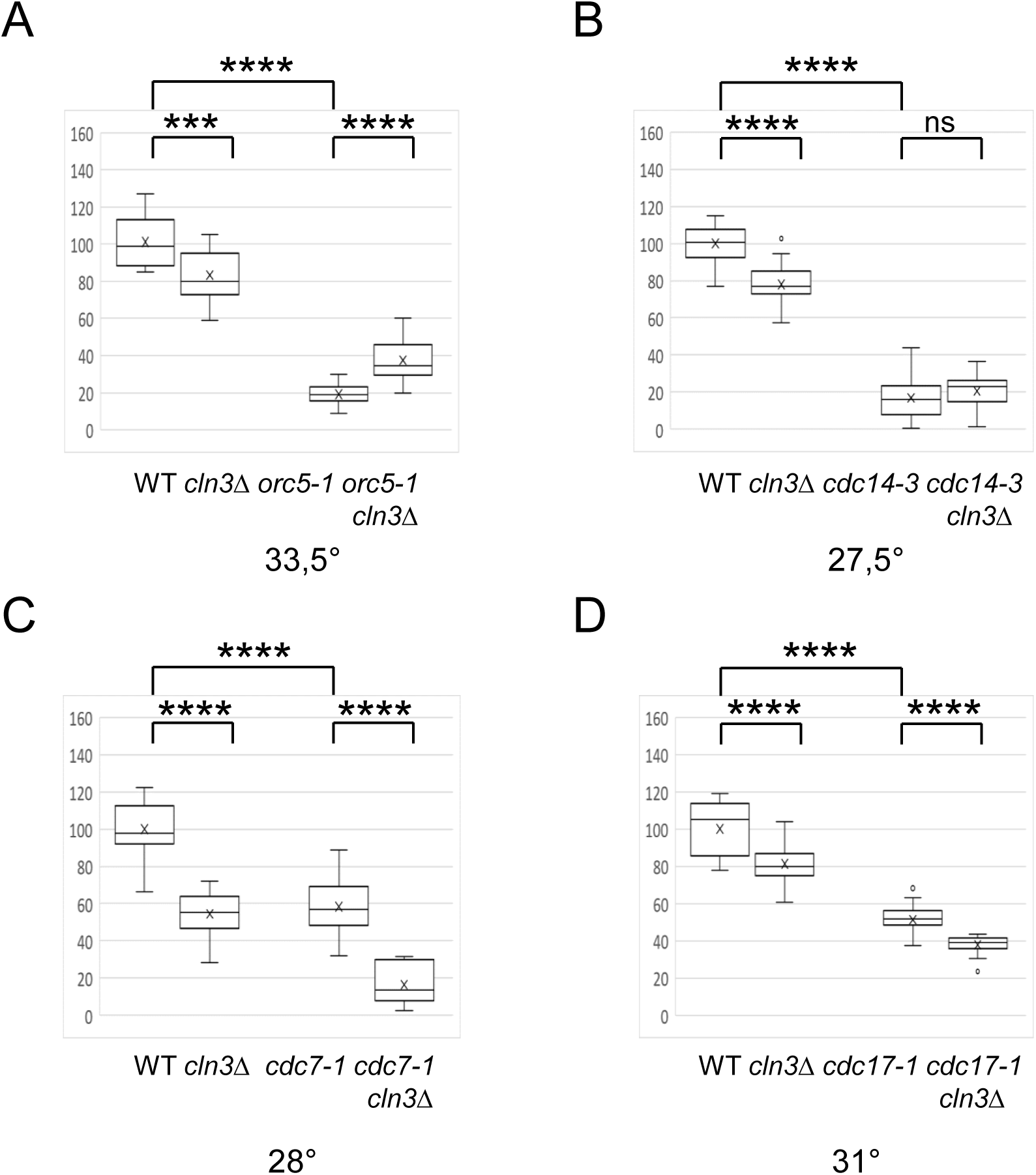
Effect of *cln3Δ* on *orc5-1*, *cdc14-3*, *cdc7-1* and *cdc17-1* spore colony size. Analysis of the meiotic products of the *ORC5/orc5-1 CNL3/cln3Δ* (A), *CDC14/cdc14-3 CNL3/cln3Δ* (B), *CDC7/cdc7-1 CNL3/cln3Δ* (C) and *CDC17/cdc17-1 CNL3/cln3Δ* (D) diploids. The box plots correspond to sizes of the spore-derived colonies grown 2-3 days at the indicated temperature according to the spore genotype. Sizes were normalized by setting the mean size of the WT at 100. P-values in a one-way ANOVA test are indicated : ns *p* > 0.05, * *p* < 0.05, ** *p* < 0.01, *** *p* < 0.001, **** *p* < 0.0001.

To confirm the idea that G1 elongation could be responsible for the positive effect of an extra chromosome, we tested the effect of Cln3 loss on *cdc14-3*, another mutation whose effect was counterbalanced by the presence of an extra chromosome (Fig 6). Unlike the wild type, no negative effect of Cln3 loss was observed on colony size in *cdc14-3* (Fig 7B). This similarity between the effect of an extra chromosome and *cln3Δ* was confirmed in *cdc7-1* and *cdc17-1*, where a negative effect of *cln3Δ* was observed as in the case of an extra chromosome (Fig 7C,D). Based on this correlation, other mutations were combined with *cln3Δ* (S12 Fig). A positive effect of Cln3 loss on colony size was observed for the *sid2-21* and the *mcm3-1* mutations. This was expected as Sid2 (Cdt1) is involved in MCM complex loading like Cdc6 and Mcm3 is one subunit of the MCM complex. In the case of *scc1-73*, a negative effect of Cln3 loss on colony size was observed, in agreement with the negative effect of CFVII for other SCC mutants (S9 Fig).

The effect of CFVII in the absence of Cln3 was studied in the case of *orc5-1* (Fig 8). In the wild type, the negative effects of *cln3Δ* and CFVII were additive. This additivity was reminiscent of that observed when CFIII and CFVII were together (S8 Fig) and confirmed the similarities in the way *cln3Δ* and an extra chromosome affect the G1 phase [27]. In *orc5-1*, no additional positive effect of CFVII was observed when Cln3 was missing. One interpretation is that the G1 phase extension due to the loss of Cln3 limits the positive effect of CFVII. This reinforces the idea that the positive effect of CFVII on *orc5-1* was achieved through the extension of G1.

**Fig 8.**
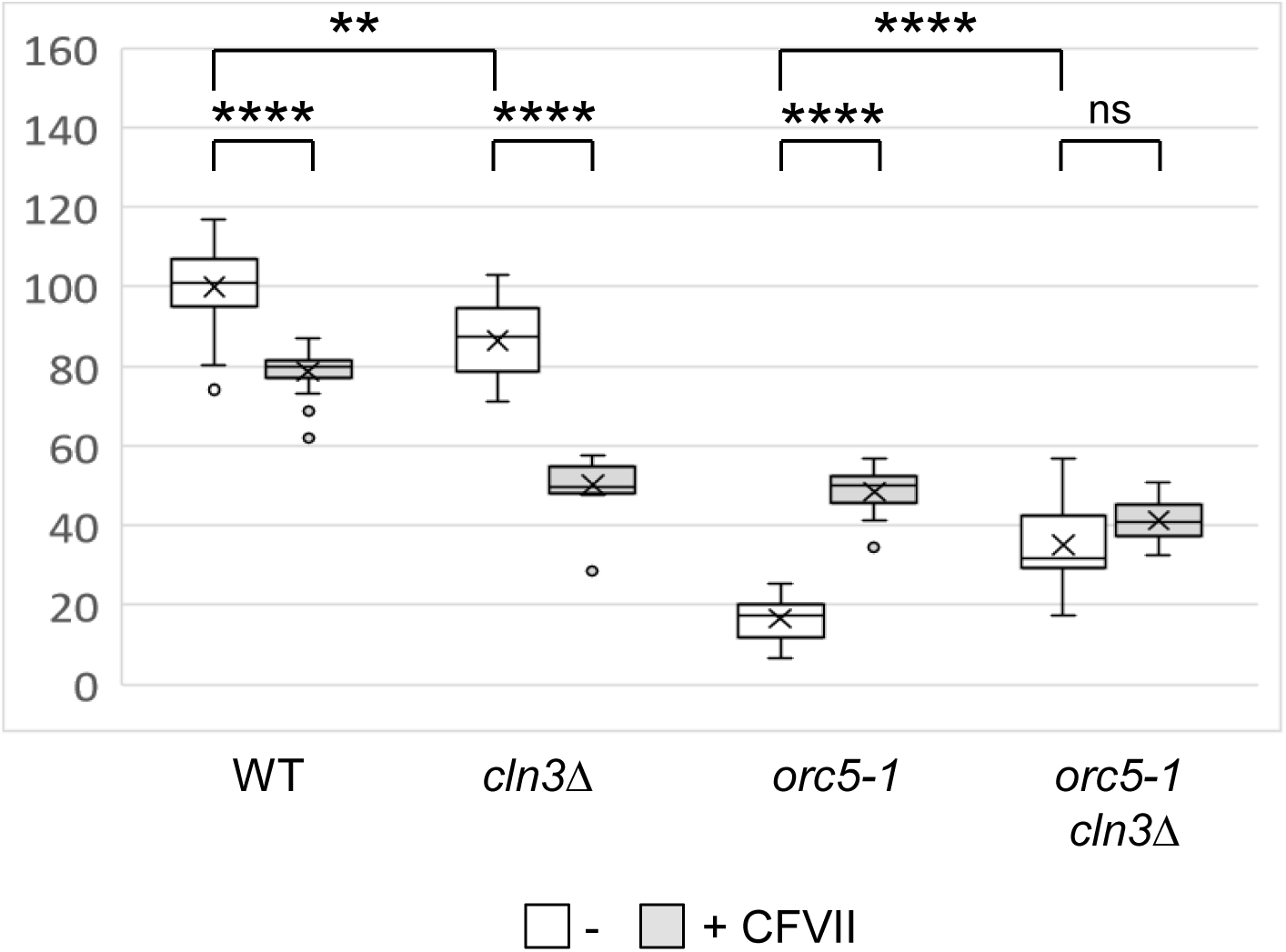
Combined effects of CFVII and *cln3Δ* on spore colony size. Analysis of the meiotic products of the *ORC5/orc5-1 CLN3/cln3Δ* diploid with CFVII. The box plots correspond to sizes of the spore-derived colonies grown 2,5 days at 33,5° on YPD according to the spore genotype and the absence/presence of CFVII. Sizes were normalized by setting the mean size of the WT without CFVII at 100. P-values in a one-way ANOVA test are indicated : ns *p* > 0.05, * *p* < 0.05, ** *p* < 0.01, *** *p* < 0.001, **** *p* < 0.0001.

### The extra chromosome limits the activation of the mitotic checkpoint

How can the lengthening of G1 shorten the total duration of the cell cycle? Defective initiation in ORC or MCM mutant cells leads to slower, incomplete replication and G2/M arrest imposed in part by the Mad2-dependent mitotic checkpoint [21,33,34]. Consequently, cell cycle arrest in response to *orc2-1*, *orc5-1*, *mcm2-1* or *mcm3-1* mutations is attenuated by the loss of Mad2. Therefore, the absence of Mad2 should have a positive effect on mutant colony size. We indeed observed that *mad2Δ* had a positive effect on *orc5-1* above a certain incubation temperature, as was the case with an extra chromosome (S13A Fig). As the loss of Mad2 increases the rate of chromosome mis-segregation [35], this positive effect could come from the acquisition of an extra chromosome. However inactivation of Chl1, also associated with increased chromosome loss [36], gave the opposite result, with a strong negative effect in *orc5-1* at 25°C, and synthetic lethality at 33.5°C (S13B Fig). Thus, the positive effect of *mad2Δ* was probably not a consequence of a defect in chromosome segregation, but rather of the reduction in cell cycle length due to attenuation of the G2/M checkpoint. This was consistent with the fact that the absence of Mad2 limits G2/M accumulation in *orc5-1* cells [21].

The decrease in G2/M accumulation in *orc5-1* cells observed in the presence of an extra chromosome (Fig 4B) suggests that the mitotic checkpoint was no longer activated. One prediction was that the loss of Mad2 should no longer have any effect in the presence of CFVII. This was indeed what has been observed, as in the presence of CFVII, the size of *orc5-1 mad2Δ* colonies was no greater than that of *orc5-1* colonies (Fig 9A). This lack of effect of *mad2Δ* suggested that attenuation of the G2/M checkpoint added nothing when G1 was lengthened in the presence of CFVII. Similarly, no additivity was observed between *mad2Δ* and *cln3Δ* mutations with regard to *orc5-1* colony size (Fig 9B).

**Fig 9.**
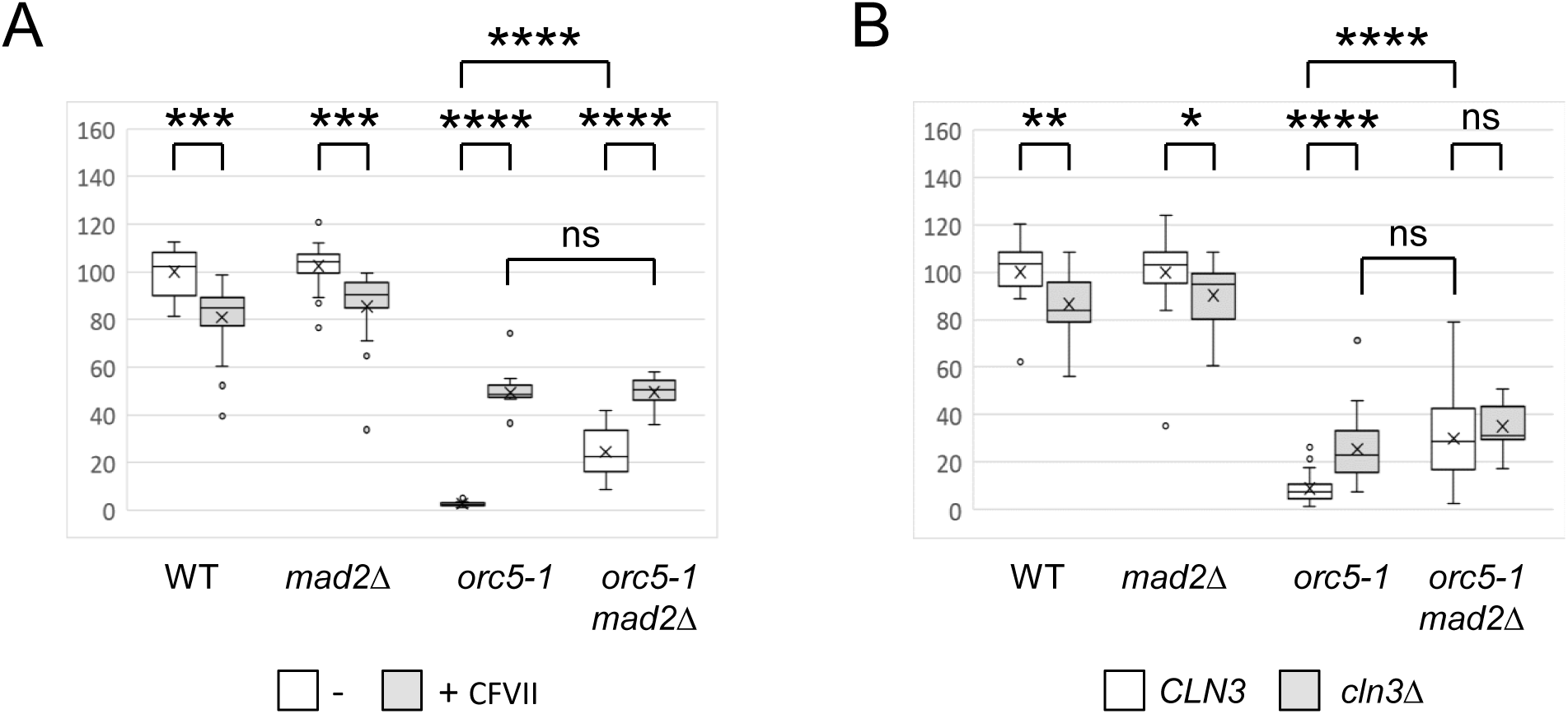
Combined effects of CFVII, *mad2Δ* and *cln3Δ* on *orc5-1* colony size. (A) Analysis of the meiotic products of the *ORC5/orc5-1 MAD2/mad2Δ* diploid with CFVII. The box plots correspond to sizes of the spore-derived colonies grown 3 days at 34° according to the spore genotype. Sizes were normalized by setting the mean size of the WT without CFVII at 100. P-values in a one-way ANOVA test are indicated : ns *p* > 0.05, * *p* < 0.05, ** *p* < 0.01, *** *p* < 0.001, **** *p* < 0.0001. (B) Same analysis as in (A) with the meiotic products of the *ORC5/orc5-1 MAD2/mad2Δ CLN3/cln3Δ* diploid. Sizes were normalized by setting the mean of the WT values at 100.

## Discussion

The work presented here was initiated on the basis of a serendipitous observation, i.e. the presence of a second copy of chromosome III in *orc5-1 set1Δ* haploid cells. This initial observation led us to investigate the reasons behind the spontaneous chromosome III disomy in this genetic context. Our results clearly show that an extra chromosome III was associated with a gain in fitness for *orc5-1 set1Δ* cells, assessed by the increased growth of spore-derived colonies, contrasting with the loss of fitness observed in the case of wild-type cells. This was observed in two different genetic backgrounds (SK1 and W303), demonstrating its general nature. Together with the fact that no increase in chromosomal instability can be detected in *orc5-1 set1Δ* (S14 Fig), this strongly suggests that the presence of an extra chromosome III has been positively selected.

The limited negative effect of chromosome III disomy that we observed in wild-type W303 cells was in apparent contradiction with previous work in which an extra chromosome III was associated with lethality in this background [37]. This difference may be linked to the way in which a second chromosome III was introduced in W303 cells : either abruptly, in one step [37], or progressively, through successive backcrosses (our work). We can assume that under the latter conditions, cells with a W303 background get the time to adapt to the negative consequences attached to the presence of an extra chromosome III.

In previous studies, the selection of an extra chromosome has been in most cases linked to an increase in the level of expression of specific genes that enable adaptation to a particular environmental or genetic stress [38]. In the case of *orc5-1 set1Δ*, it could be assumed that the chronic replicative stress that exists in this mutant [21] was alleviated by the overexpression of one or several chromosome III genes. Inactivation of two candidate genes, *MRC1* and *DCC1*, on the left arm of chromosome III had no impact on the positive effect of III disomy on *orc5-1 set1Δ* (S4 fig). Together with the fact that a fragment of chromosome VII could recapitulate the effect of chromosome III (S5 Fig), this favors a general consequence of aneuploidy independent of the genes on the duplicated chromosome. For this reason, we believe that duplication of a chromosome other than III could also be selected in *orc5-1 set1Δ* context.

By increasing the temperature to which the spore-derived colonies were grown, we then discovered that the positive effect of an extra chromosome acted on the *orc5-1* mutation, independently of the *set1Δ* mutation (Fig 3). It therefore appears that this positive effect on *orc5-1* was only visible when the *orc5-1* mutation had sufficient impact on cell physiology, i.e. either above a certain temperature, or when associated with a sensitizing mutation such as *set1Δ*.

While an extra chromosome had a negative effect for mutations affecting proteins involved in DNA replication (Cdc7, Cdc17) and cohesion (Scc2, Smc1, Smc3), it showed a positive effect for mutations other than *orc5-1*. These included *orc2-1*, which, like *orc5-1*, affects an ORC subunit, *cdc6-4*, required for the pre-RC assembly, but also *cdc14-3* and *cdc15-2*, affecting mitotic exit. This raised the question of a possible functional link between these different proteins. Importantly, beyond its established function in mitotic exit, Cdc14 plays an important role in origin licensing, with pre-RC components such as Orc2 and Cdc6 being Cdc14 substrates [39]. The role of Cdc14 as a key element in the assembly of pre-RCs at the end of mitosis, by stabilizing Cdc6 through dephosphorylation, has recently been confirmed [40]. Furthermore, the impact of the *cdc15-2* mutation on origin licensing is comparable to that of the *cdc14-3* mutation [39]. Thus, it turns out that among the mutations tested, those for which a positive effect of the extra chromosome could be observed were directly (*orc2-1*, *orc5-1, cdc6-4*) or indirectly (*cdc14-3*, *cdc15-2*) involved in origin licensing.

The gain of an extra chromosome has been shown to interfere with cell proliferation, essentially by lengthening the G1 phase [1,27]. As the G1 phase is the period of the cell cycle during which pre-RC complexes are assembled [41], we hypothesized that an extension of G1 could explain the positive effect of an extra chromosome in the presence of mutations that affect pre-RC assembly. This hypothesis is supported by: (i) the lack of a positive effect of YACs on the growth of *orc5-1* colonies (S7 Fig) that correlates with the absence of a delay in cell cycle entry in YACs-bearing strains [1,28]; (ii) the opposite effects of eliminating Cln3 (increased G1 duration), and overexpressing Cln3 (decreased G1 duration), with positive and negative effect on the growth of *orc5-1* colonies respectively (Fig 7 and S11 Fig). The parallel between the effect of an extra chromosome and the loss of Cln3 can be extrapolated to other mutations. The positive effect of *cln3Δ* in the case of *sid2-21* and *mcm3-1* suggested that G1 elongation benefited other mutations affecting pre-RC components.

How can the increase in G1 duration offset the impact of mutations affecting the assembly of functional pre-CRs? These mutations, which render the corresponding proteins thermosensitive, can affect their function and/or quantity. For example, the stability and DNA binding activity of the ORC complex is strongly reduced or eliminated in *orc5-1* and *orc2-*1 cells at the non-permissive temperature [34,42]. At a semi-permissive temperature (from 30°C to 34°C), we can assume that the stability and activity of the ORC complex is only partially affected. Under these conditions, a longer G1 phase could compensate for the weakened binding of the ORC complex at some replication origins. Even at a permissive temperature, only a subset of origins is activated in *orc5-1* [43], which is reflected by the mild growth defect of *orc5-1* colonies that we observe at 25°C and could explain the absence of negative effect of an extra chromosome at this temperature (Fig 1). As not all origins bind ORC with the same efficiency [44], the positive effect of extending G1 could relate specifically to those origins which are the least efficient at binding ORC. Alternatively, a longer G1 phase may allow Orc proteins to accumulate to a higher level, leading to more licensed origins at the start of the S phase.

Whatever the mechanism at play, it appears that the benefit of an extension of the G1 phase exceeds the costs of an extra chromosome in mutants with perturbed pre-RC assembly, with an effect on colony growth that is the opposite of that in the wild type (see the model shown in Fig 10). The lower number of active origins resulting from a defect in pre-RC formation leads to a prolongation of the S phase associated with some replicative stress [45], that activate the mitotic checkpoint [46]. We propose that the extra time given by the lengthening of the G1 phase in the presence of a supernumerary chromosome provides an opportunity to compensate for the defect in pre-RC formation, leading to an increase in the number of licensed origins. The consequence is a reduction in S-phase duration, and hence in replicative stress and mitotic checkpoint activation, as supported by the absence of effect of Mad2 loss in the presence of CFVII (Fig. 9A). The reduction in time spent in the S and G2/M phases would outweigh the lengthening of the G1 phase, resulting in a shortening of the total cell cycle duration. The fact that an extra chromosome had a negative effect in the absence of Sic1 (Fig 6) indirectly corroborates our model. Sic1 is an inhibitor of S-phase CDKs and the partial defect in origin licensing of *sic1Δ* cells is due to early activation of S CDKs, whose activity is known to interfere with pre-RC assembly [47]. Extending the duration of the G1 phase, while S CDKs are active in the absence of Sic1, should therefore not increase the number of pre-RC complexes formed in this mutant.

**Fig 10.**
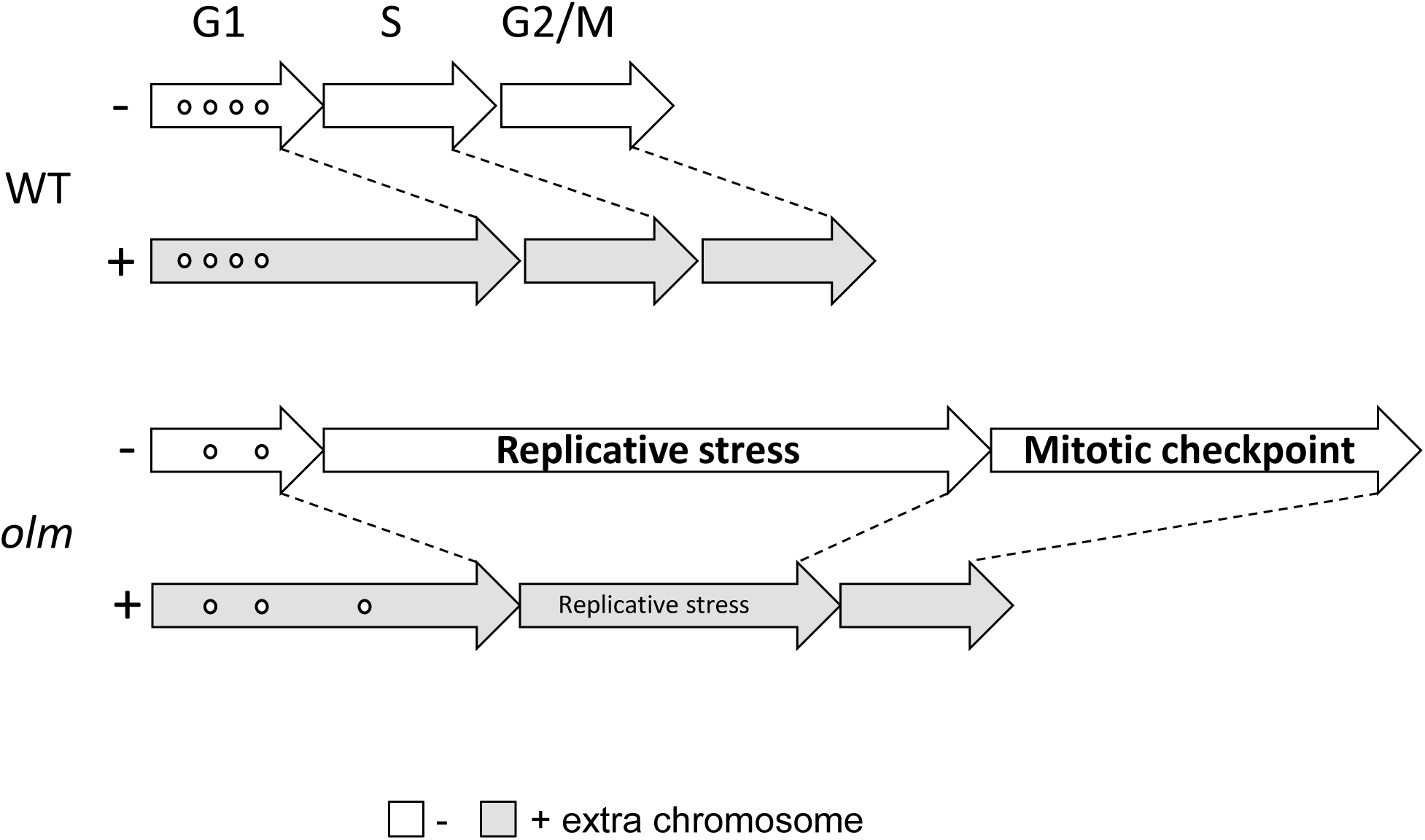
A model to explain the positive effect of an extra chromosome in a mutant affecting origin licensing. The cell cycle is represented by three successive arrows corresponding to the G1, S, G2/M phases. The small circles in the G1 arrow symbolize functional pre-RC complexes. In the wild type, the presence of an extra chromosome (grey arrows) leads to a lengthening of the G1 phase, resulting in a slower growth rate. In an origin licensing mutant (olm), the number of functional pre-RCs is reduced. This results in a lengthening of the S phase, accompanied by replicative stress which may activate the mitotic checkpoint, leading to a lengthening of the G2/M transition. The lengthening of the G1 phase, linked to the presence of an extra chromosome, can give time to increase the number of functional pre-CRs. This would result in reduced S-phase duration, replicative stress and mitotic checkpoint activation, thus decreasing total cell cycle duration and increasing growth rate.

To observe the positive effect of an additional chromosome on the fitness of a pre-RC mutant, certain conditions must be met, as illustrated in the case of *orc5-1*. First of all, the amount of additional genetic material must not be too great, as shown by the negative effect associated with the presence of large chromosome XII (S6 Fig) or the combined presence of CFIII and CFVII (S8 Fig). In such a situation, the costs associated with overexpressing too many genes seem to outweigh the benefits of extending the G1 phase. Secondly, certain environmental parameters come into play. The growth temperature must be adapted to the level of thermosensitivity of the mutation in question. Equally important is the culture medium used, which itself has an effect on the duration of the G1 phase. It appears that the faster the growth, the shorter the G1 phase, and the greater the positive effect of an extra chromosome.

Our observation of a positive effect of an extra chromosome in certain genetic contexts in *Saccharomyces cerevisiae* can potentially be extrapolated to mammalian cells, particularly cancer cells in which numerous oncogenic mutations generally coexist with aneuploidy [48]. Importantly, oncogene activation has been shown to constitute an important source of replication stress that likely drives cancer development [49]. Consequently, cancer cells must cope with replication stress in order to proliferate [50]. In line with the findings of this work, the fact that an extra copy of a chromosome leads to G1 phase elongation in mammalian cells [1,5,7], and the evidence linking oncogene-induced replication stress to defects in origin licensing [51], we can postulate that one reason among others for aneuploidy selection in cancer cells is its ability to mitigate replication stress by increasing the G1 phase duration and, consequently, the number of licensed origins.

## Materials and Methods

### Yeast strains and media

The strains of *S. cerevisiae* used in this study are listed in S1 Table. The *RER1* gene located near the centromere of chromosome III was replaced by either the *hph*MX marker, conferring resistance to hygromycin B, or the *nat*MX marker, conferring resistance to nourseothricin. The presence of chromosome III disomy in haploid cells, which can be inferred from the presence of both markers, was confirmed by pulsed-field gel electrophoresis (PFGE) as described [22].

YPD medium (1% [w/v] yeast extract, 2% [w/v] peptone, 2% [w/v] glucose) was used as the standard medium for general culture. For sporulation induction, cells were directly striked from YPD plates on 1% potassium acetate sporulation medium plates.

Spores were isolated on YPD medium or on synthetic minimal SD medium (0.17% [w/v] yeast nitrogen base without amino acids and ammonium sulfate, 0.5% [w/v] ammonium sulfate, 2% [w/v] glucose supplemented with required amino acids). For experiments in which the *GAL1* promoter was used to drive *CLN3* expression, spores were isolated on YPGal (1% [w/v] yeast extract, 2% [w/v] peptone, 2% [w/v] galactose).

### Spore colony size measurement

Tetrad dissection was performed using the MSM 400 dissection microscope (Singer Instrument Company) and isolated spores were incubated for two to four days depending on the medium and incubation temperature. JPEG files of dissection plates were analyzed with the image-processing software ImageJ [23] to measure the area of each spore-derived colony. Genotypes of individual spore colonies were determined by replica plating to selective media or temperature.

In dissections where CFIII and CFVII were combined, spores containing either chromosomal fragment were genetically indistinguishable, as the same genetic markers were present on each. These spores could be differentiated from spores containing both CFIII and CFVII on the basis of the distribution of genetic markers within each tetrad. When the genetic markers were present in all four spores of a tetrad, this meant that each spore had either CFIII or CFVII. When the genetic markers were present in only two spores of a tetrad, this meant that they possessed both CFIII and CFVII. When the genetic markers were present in three spores of a tetrad, this meant that two of them had either CFIII or CFVII, and the third had both CFIII and CFVII. In the latter case, the size of the colonies made it possible to distinguish between the two categories of spores.

### Analysis of spore colony cells by flow cytometry

Cells from spore-derived colonies were harvested directly from YPD plates and resuspended in PBS, then fixed in 70% ethanol at room temperature. After rehydration in PBS and brief sonication to remove cell clumps, samples were incubated for 2 h with RNase A (1 mg/ml) at 42°C, then for 30 min with proteinase K (2 mg/ml) at 50°C. After washing in PBS, cells were resuspended in 1μM Sytox Green, and DNA content was determined by cytofluorimetry with a Becton Dickinson Accuri C6. Forward scatter measurement (FSC) was used to determine cell size on the same samples.

## Acknowledgments

We thank Valérie Garcia, Christelle Cayrou, Marie-Noëlle Simon, Dmitri Churikov, Karel Naiman and Eric Bailly for critical reading of the manuscript; Ndieme Cisse, Loïc Vuillermet, Saliha Lagsier and Mariam Slimani for their contribution in this work during their training course in the laboratory; Michael Weinreich for the *orc5-1*, *cdc6-4* and *cdc17-1* strains; Maitreya Dunham for strains carrying YAC-2 and YAC-3; Etienne Schwob for the *sid2-21*, *mcm3-1*, *cdc14-3*, *cdc15-2* and *sic1Δ* strains; Doug Kellog for the *cln3Δ* and *GAL1-CLN3* strains; Kim Nasmyth for the *scc1-73*, *smc1-259*, *smc3-42* and *scc2-4* strains. This work was supported by the Ligue Nationale Contre le Cancer (LNCC; Equipe labellisée).

## Figure legends

**S1 Fig.**
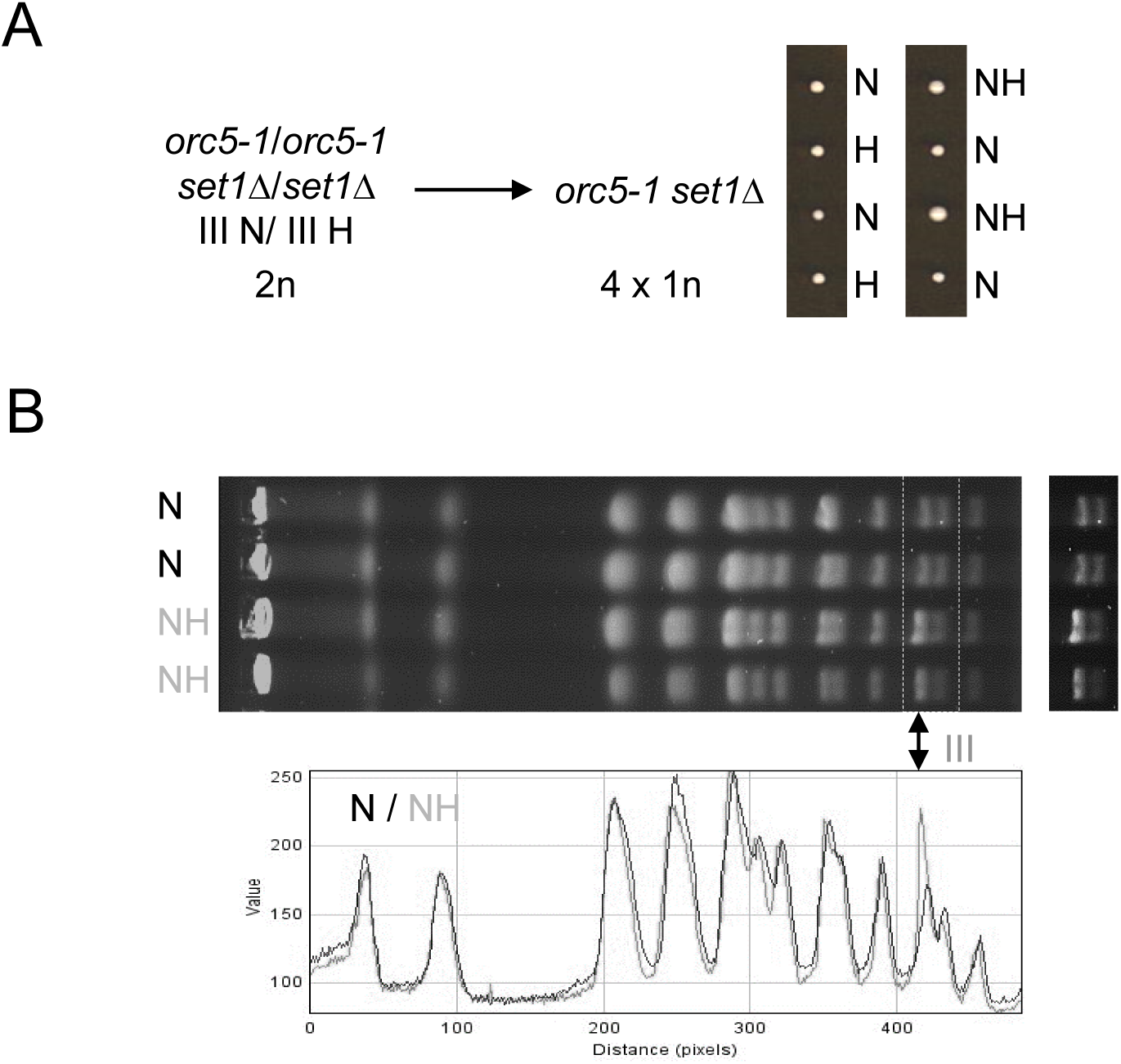
Demonstration of a chromosome III disomy in *orc5-1 set1Δ*. (A) A *orc5-1/orc5-1 set1Δ/set1Δ* diploid (2n) with chromosomes III bearing either the *Nat*MX or the *Hph*MX centromeric markers (III N/ III H) was sporulated. The colonies shown on the right correspond to the four haploid spores (1n) isolated from two tetrads. N and H: genetic markers *Nat*MX and *Hph*MX. (B) Top : chromosomes from cells derived from four spores of the same N/N/NH/NH tetrad were separated by PFGE. On the right is a more contrasted exposure of the area including the chromosome III. Below : superimposition of quantitative profiles obtained for each spore category, either with a single marker (N, black) or two markers (NH, grey).

**S2 Fig.**
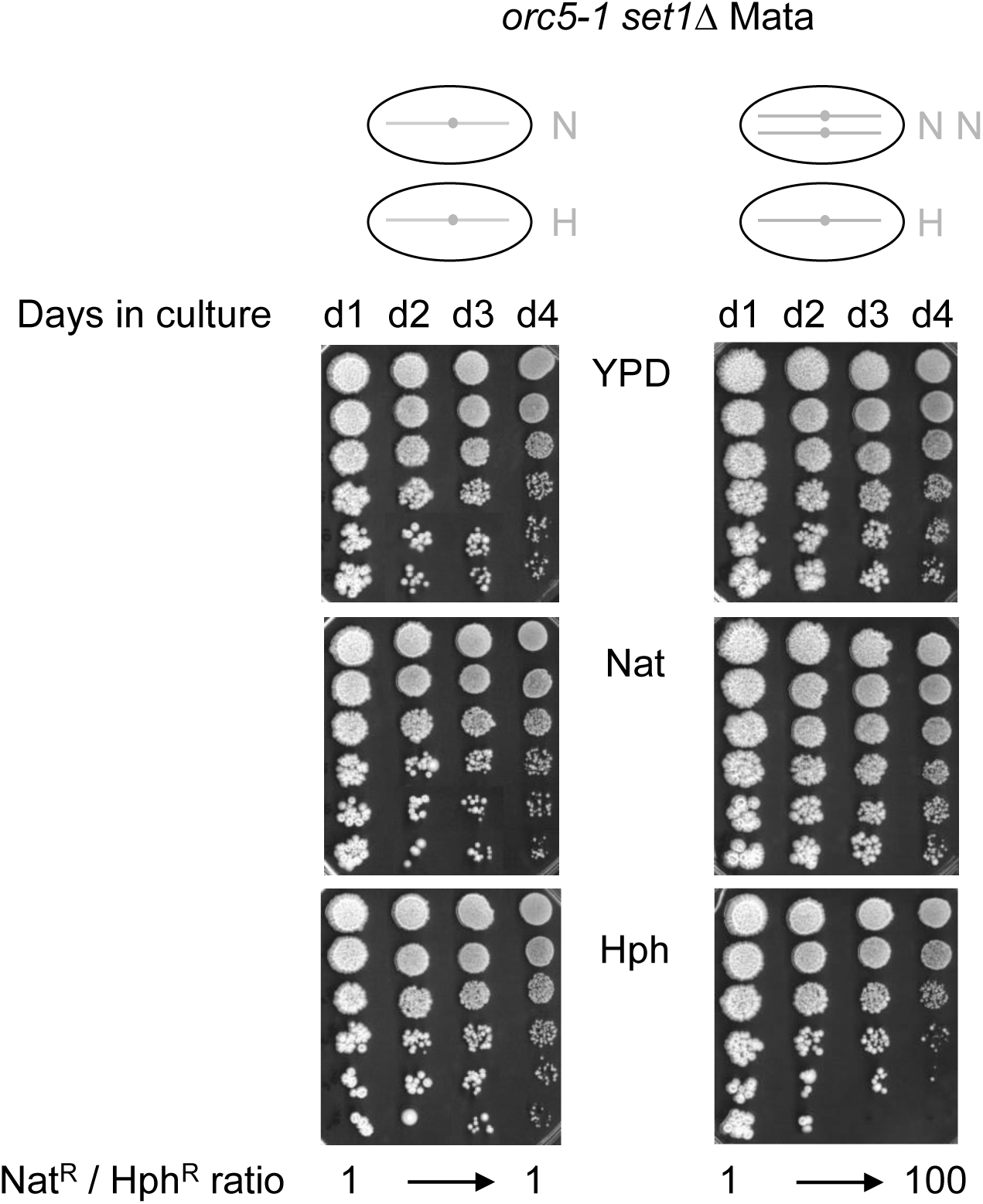
Coculture of *orc5-1 set1Δ* cells with one or two chromosome III. Equal numbers of *orc5-1 set1Δ* Mata cells with one *Hph*MX-tagged chromosome III and *orc5-1 set1Δ* Mata cells with one (left) or two (right) *Nat*MX-tagged chromosomes III were mixed and grown at 25° in YPD liquid culture medium. At one-day intervals, ten-fold dilutions of culture aliquots were spotted onto the same YPD, YPD + nourseothricin (Nat) and YPD + hygromycin (Hph) plates prior to incubation at 25°. The ratio of Nat^R^ to Hph^R^ cells is indicated at the bottom.

**S3 Fig.**
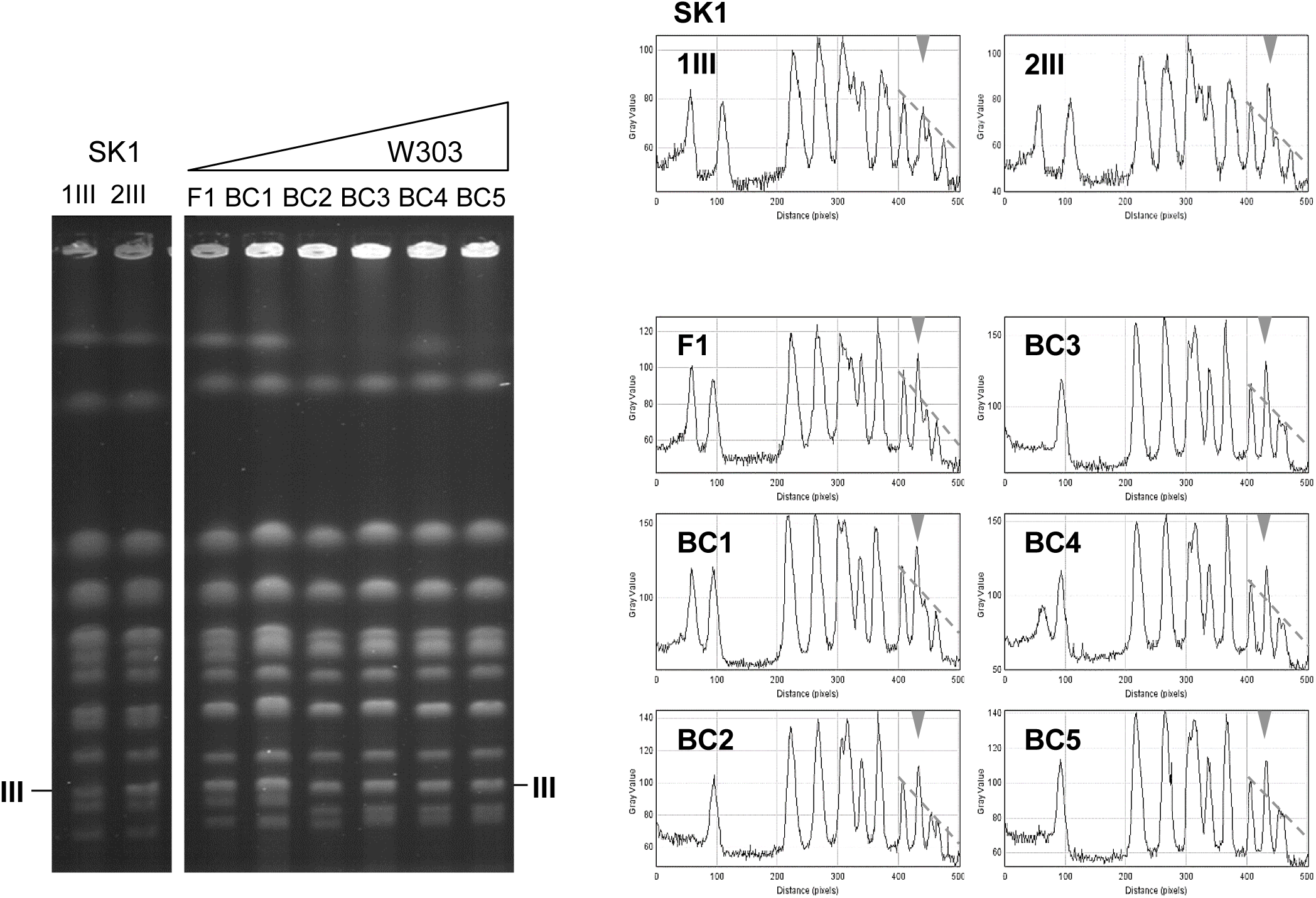
Efficient transmission of the extra copy of chromosome III during successive backcrosses. Left: the chromosomes of representative clones from each backcross, from the initial SK1/W303 hybrid (F1) to the fifth backcross (BC5), were separated by PFGE. Control: SK1 clones with one or two chromosome III. The position of chromosome III is indicated. Some differences in the chromosome size pattern exist between SK1 and W303 backgrounds. Right: quantitative profiles corresponding to each gel lane. Chromosome III is identified by an inverted grey triangle. A grey dotted line connects the peaks of the chromosome signals in the area where their height is proportional to the chromosome length.

**S4 Fig.**
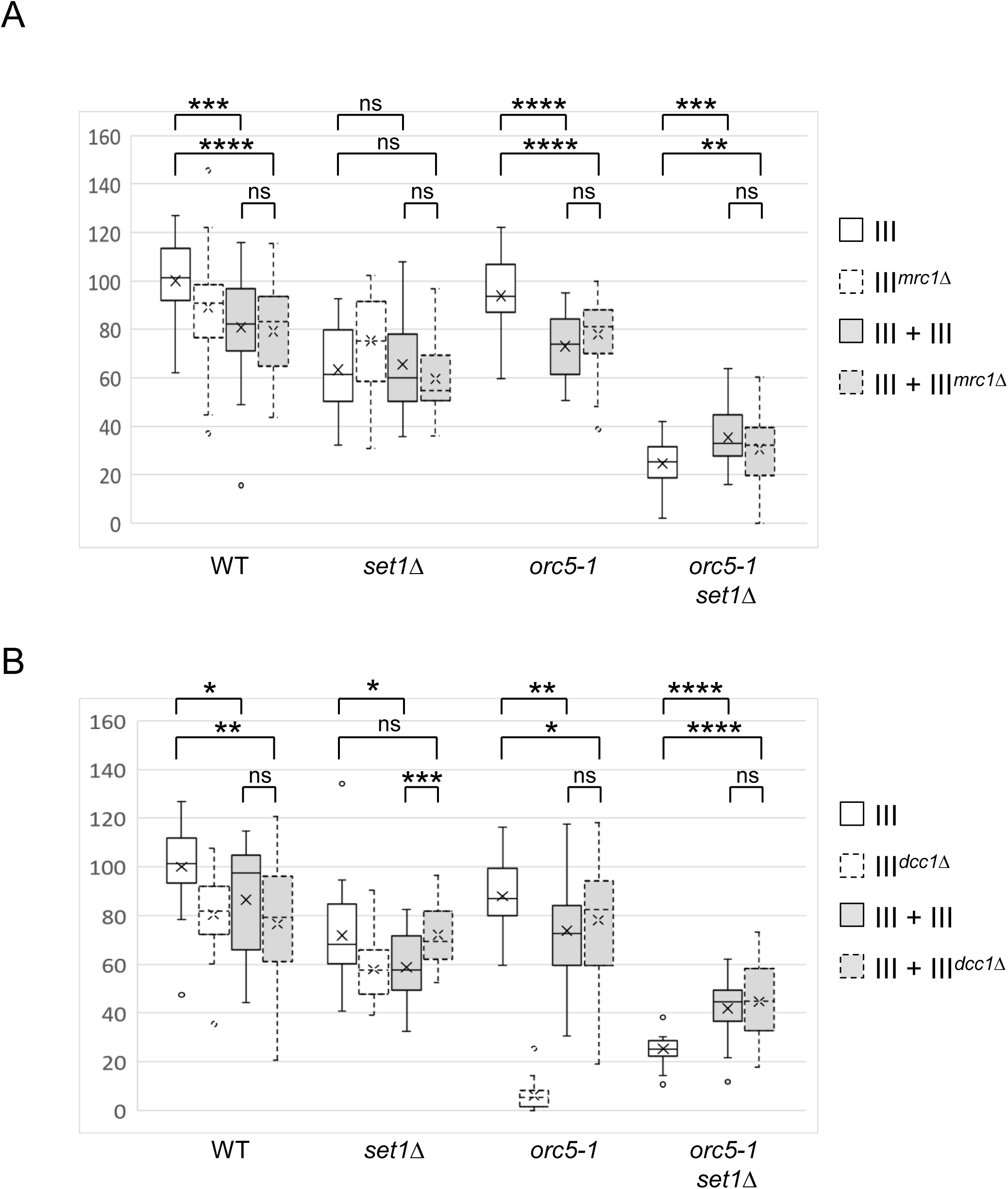
No impact of deletion of *MRC1* or *DCC1* on the effect of chromosome III disomy. (A) Analysis of the meiotic products of *ORC5/orc5-1 SET1/set1Δ* diploids with three chromosomes III, one of which carrying a deletion of the *MRC1* gene. The box plots correspond to sizes of the spore-derived colonies, grown 3 days at 25° on YPD, according to the spore genotype, the number of chromosome III and the presence of *mrc1Δ*. Sizes were normalized by setting the mean size of the WT with one III at 100. P-values in a one-way ANOVA test are indicated : ns *p* > 0.05, * *p* < 0.05, ** *p* < 0.01, *** *p* < 0.001, **** *p* < 0.0001. No boxplot: synthetic lethality. (B) Same analysis as in (A) with deletion of the *DCC1* gene.

**S5 Fig.**
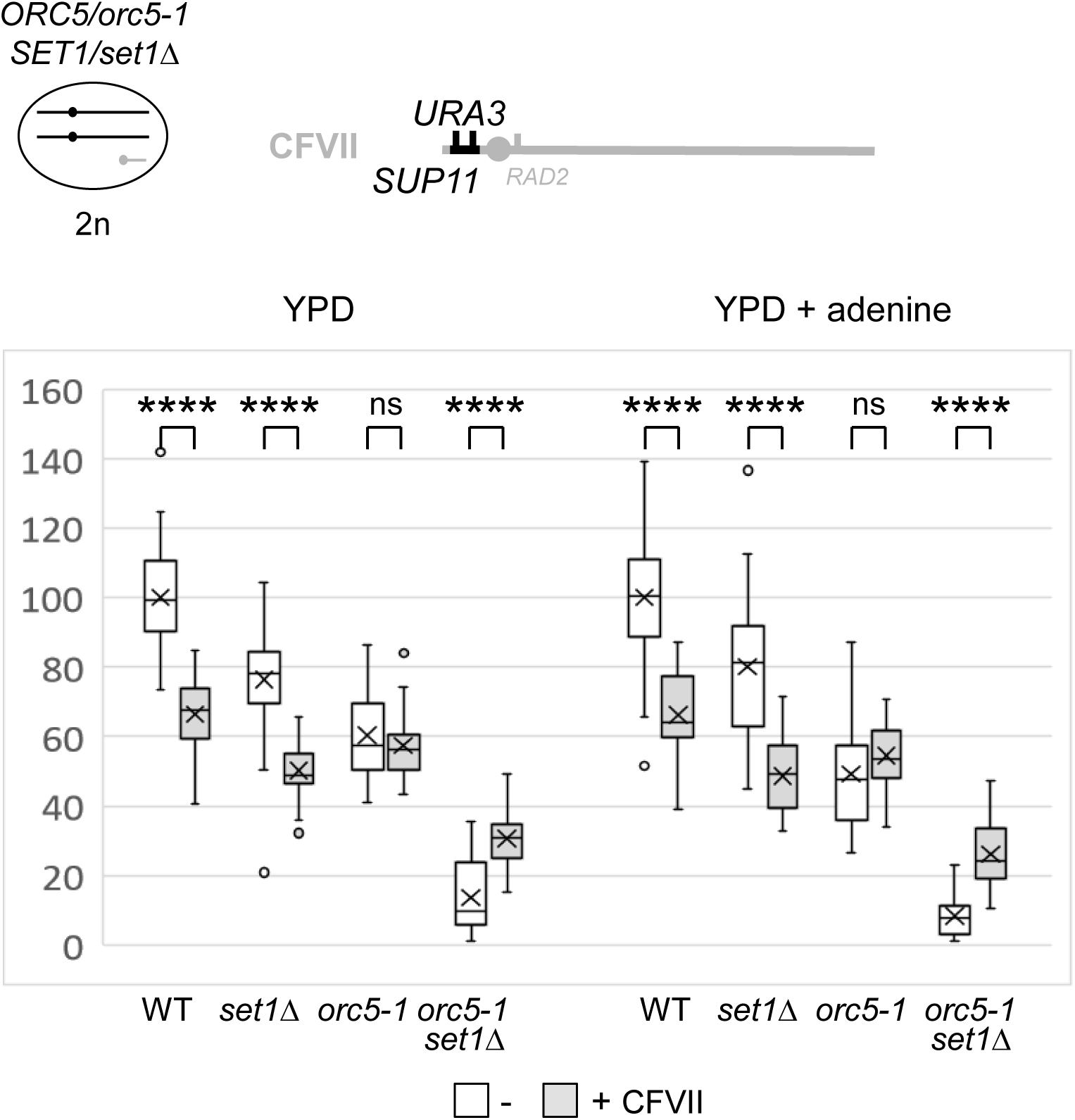
A growth advantage is associated with an extra right arm of chromosome VII in *orc5-*1 set1Δ. Top left : the *ORC5/orc5-1 SET1/set1Δ* diploid (2n) with two chromosomes VII (in black) and the additional chromosome VII fragment (CFVII in grey). Top right : schematic drawing of CFVII with its *URA3* and *SUP11*genetic markers (in black). Bottom: the box plots correspond to sizes of the spore-derived colonies, grown 3 days at 25° on standard YPD plates (left) or YPD plates supplemented with adenine (right), according to the spore genotype and the absence/presence of CFVII. Sizes were normalized by setting the mean size of the WT without CFVII at 100. P-values in a one-way ANOVA test are indicated : ns *p* > 0.05, * *p* < 0.05, ** *p* < 0.01, *** *p* < 0.001, **** *p* < 0.0001.

**S6 Fig.**
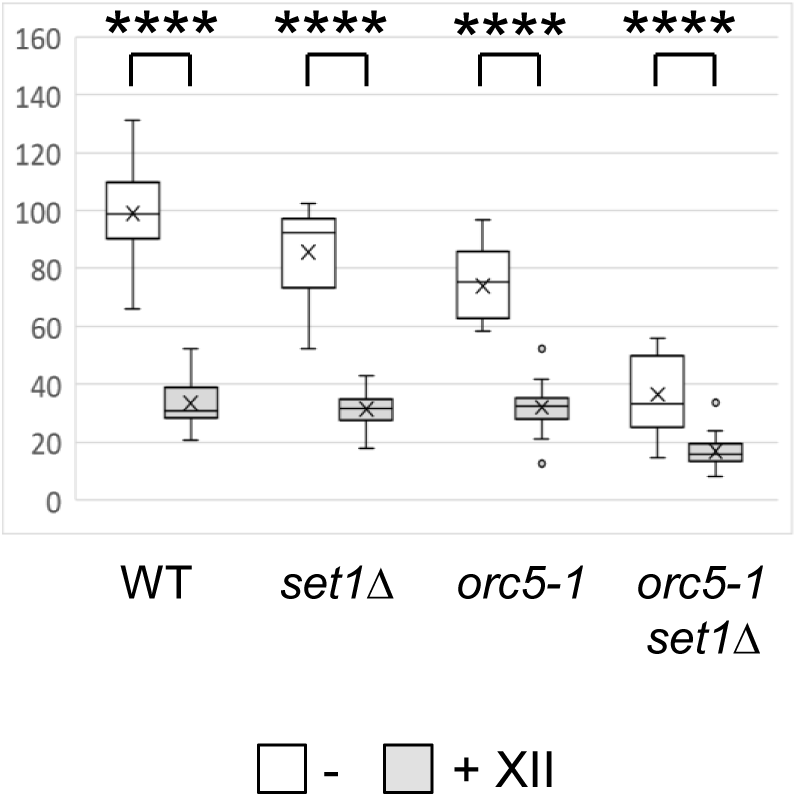
No growth advantage is associated with chromosome XII disomy in *orc5-1 set1Δ*. Analysis of the meiotic products of *ORC5/orc5-1 SET1/set1Δ* diploids with an extra chromosome XII. The box plots correspond to sizes of the spore-derived colonies grown 3 days at 25° on YPD, according to the spore genotype and the absence/presence of an extra chromosome XII. Sizes were normalized by setting the mean size of the WT without an extra chromosome XII at 100. P-values in a one-way ANOVA test are indicated : ns *p* > 0.05, * *p* < 0.05, ** *p* < 0.01, *** *p* < 0.001, **** *p* < 0.0001.

**S7 Fig.**
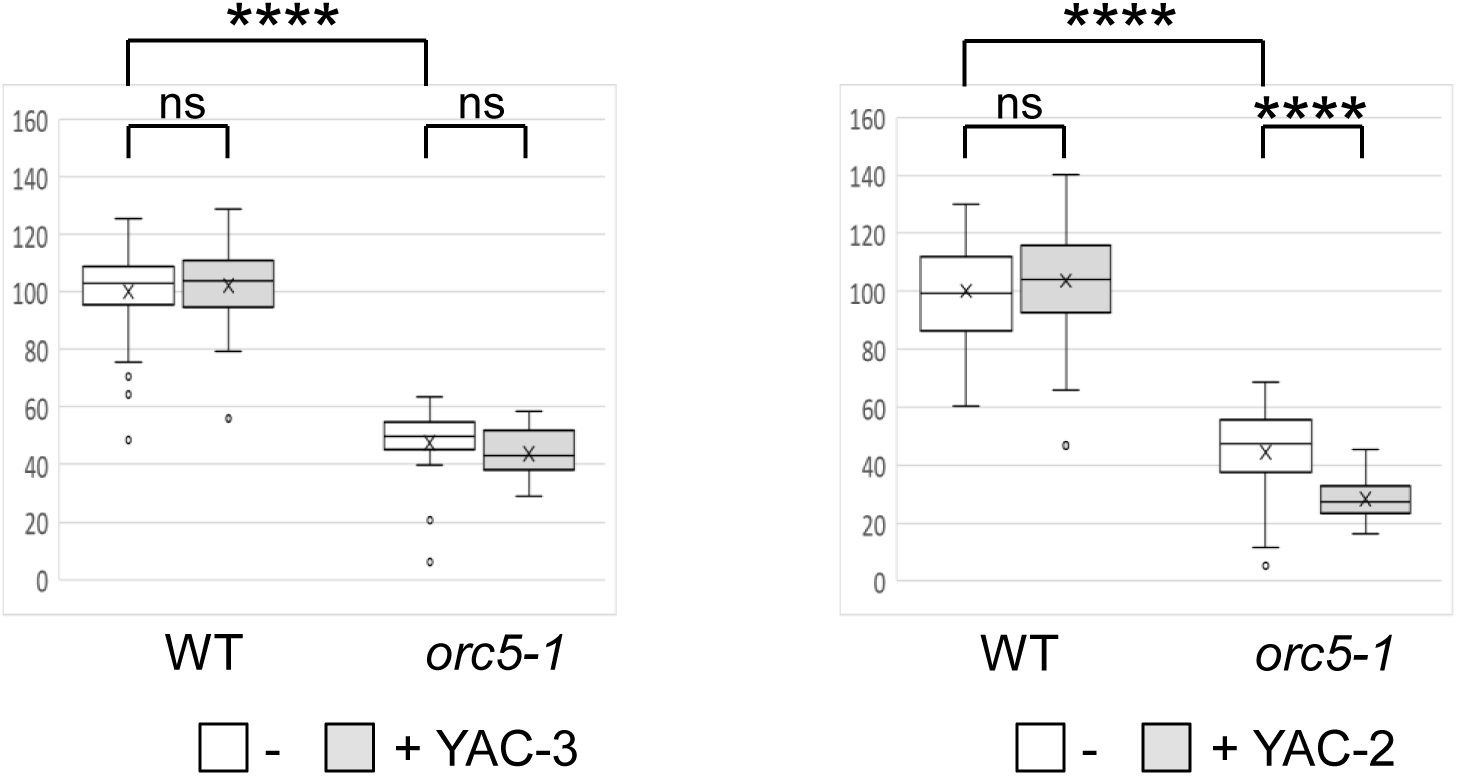
No positive effect of YAC on *orc5-1* spore colony size. Analysis of the meiotic products of heterozygous *ORC5/orc5-1* diploids with one YAC. The box plots correspond to sizes of the spore-derived colonies grown 3 days at 30° on YPD, according to the spore genotype and the absence/presence of the YAC-3 (left) or YAC-2 (right). Sizes were normalized by setting the mean size of the WT without YAC at 100. P-values in a one-way ANOVA test are indicated : ns *p* > 0.05, * *p* < 0.05, ** *p* < 0.01, *** *p* < 0.001, **** *p* < 0.0001.

**S8 Fig.**
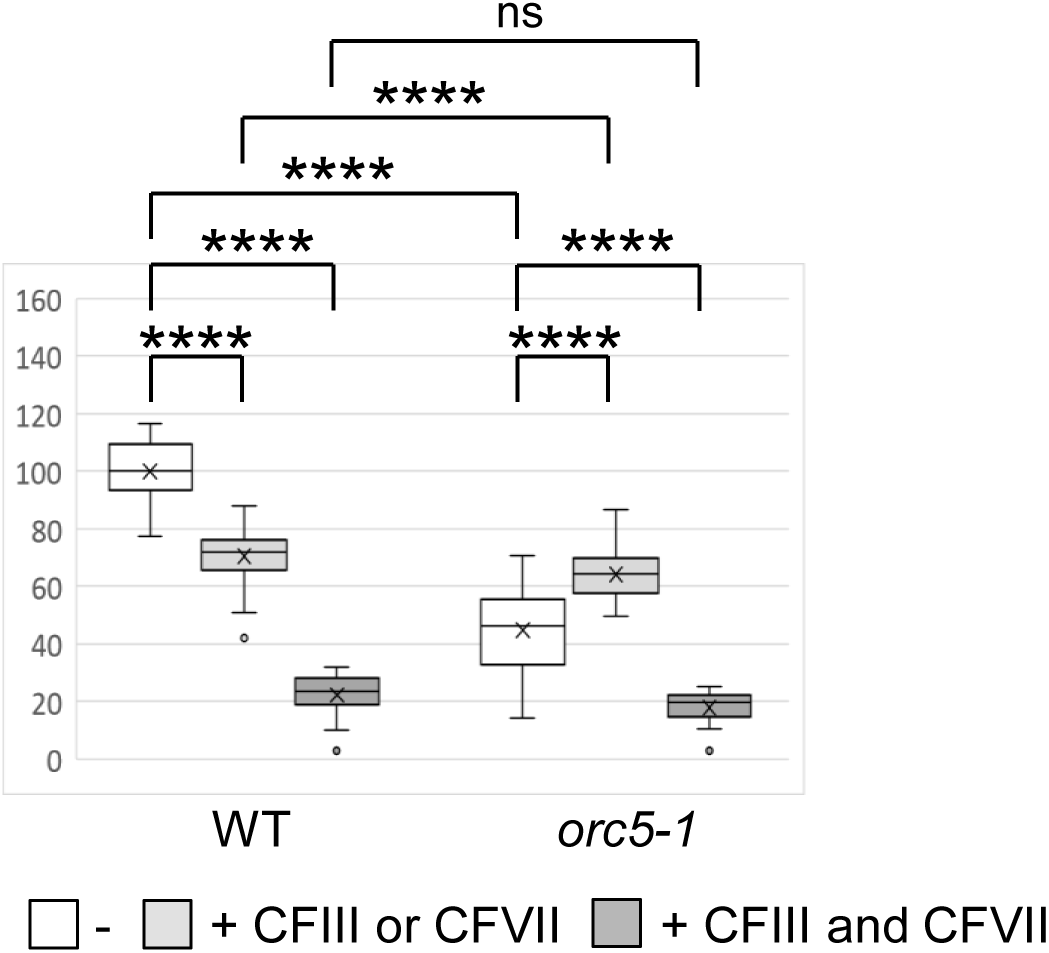
Comparing the effect of one and two chromosomal fragments on spore colony size. Analysis of the meiotic products of heterozygous *ORC5/orc5-1* diploids with CFIII and CFVII. The box plots correspond to sizes of the spore-derived colonies grown 3 days at 30° on YPD, according to the spore genotype and the presence of only one (CFIII or CFVII) or both (CFIII and CFVII) chromosome fragments. Sizes were normalized by setting the mean size of the WT without CF at 100. P-values in a one-way ANOVA test are indicated : ns *p* > 0.05, * *p* < 0.05, ** *p* < 0.01, *** *p* < 0.001, **** *p* < 0.0001.

**S9 Fig.**
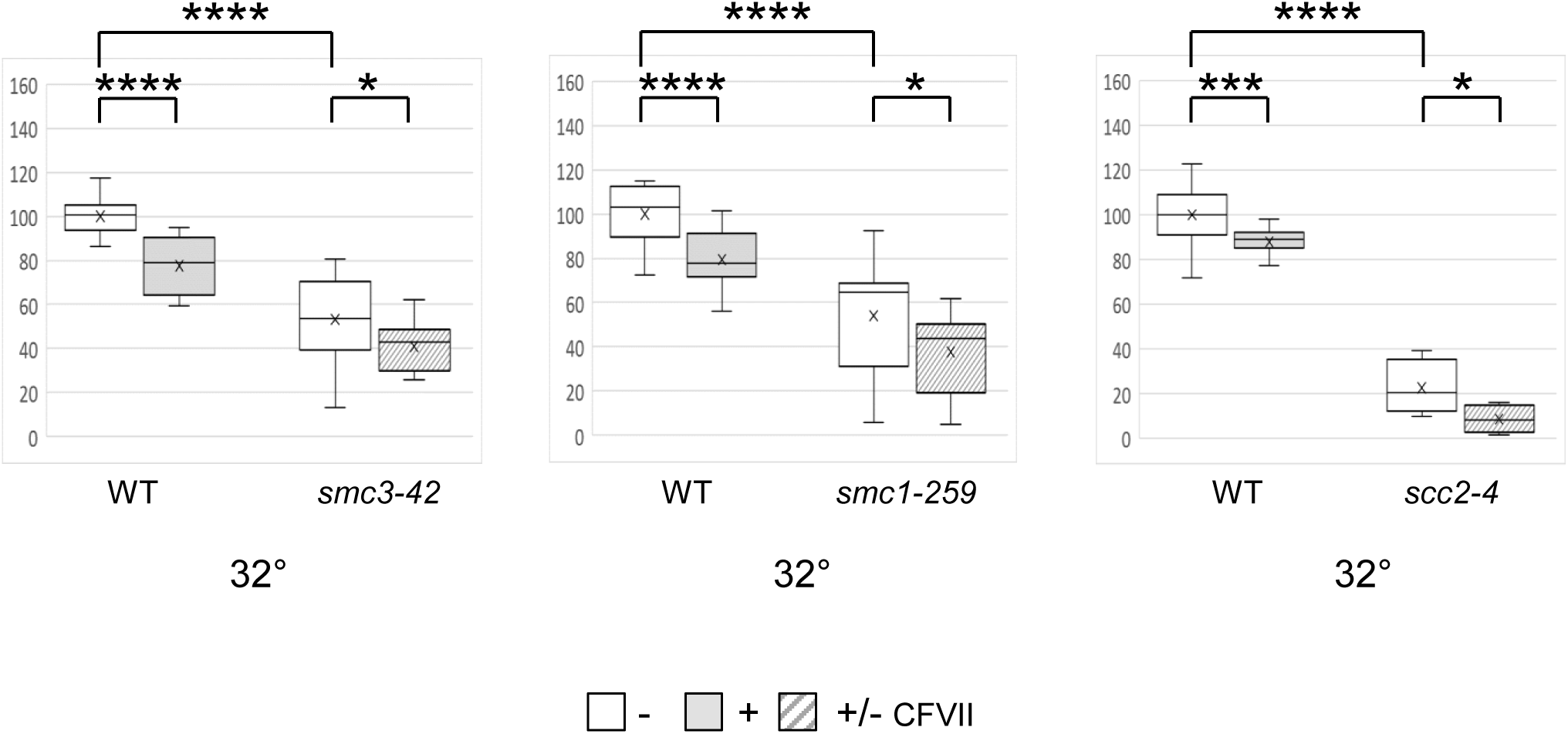
Effect of CFVII on *smc3-42*, *smc1-259* and *scc2-4* spore colony size. Analysis of the meiotic products of the *SMC3/smc3-42* (left), *SMC1/smc1-259* (middle) and *SCC2/scc2-4* (right) with CFVII. The box plots correspond to sizes of the spore-derived colonies grown 2-3 days at the indicated temperature according to the spore genotype and the absence/presence of CFVII. Stripped boxes correspond to a mix of cells with and without CFVII in mutant spore-derived colonies. Sizes were normalized by setting the mean size of the WT without CFVII at 100. P-values in a one-way ANOVA test are indicated : ns *p* > 0.05, * *p* < 0.05, ** *p* < 0.01, *** *p* < 0.001, **** *p* < 0.0001.

**S10 Fig.**
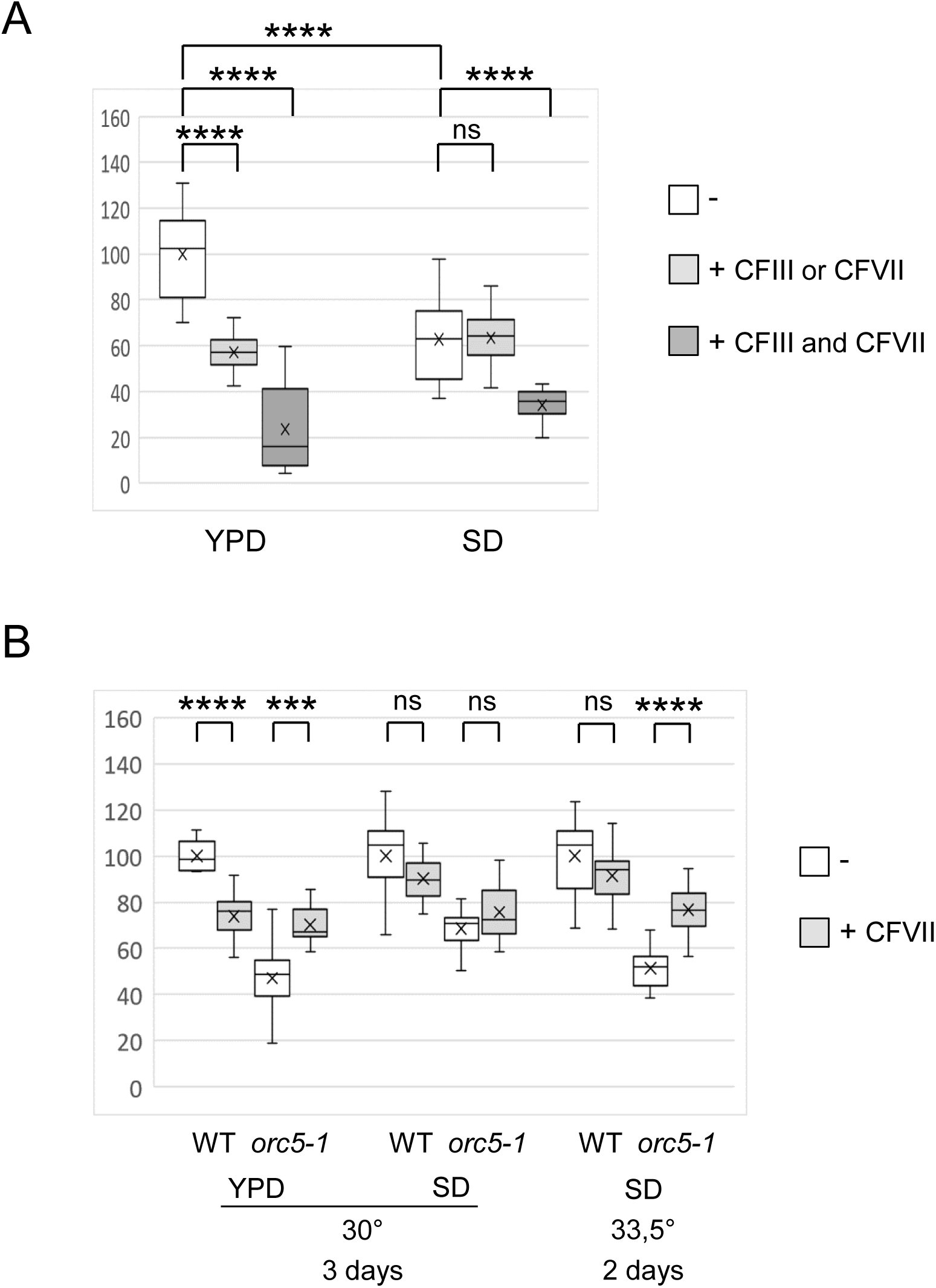
The effect of an extra chromosome depends on the growth medium. (A) Analysis of the meiotic products of the WT diploid with CFIII and CFVII. The box plots correspond to sizes of the spore-derived colonies grown 3 days (YPD) or 4 days (SD) at 25° according to the presence of only one (CFIII or CFVII) or both (CFIII and CFVII) chromosome fragments. Sizes were normalized by setting the mean size of the WT without CFVII on YPD at 100. P-values in a one-way ANOVA test are indicated : ns *p* > 0.05, * *p* < 0.05, ** *p* < 0.01, *** *p* < 0.001, **** *p* < 0.0001. (B) Same analysis as in (A) with the meiotic products of *ORC5/orc5-1* diploids with CFVII. The box plots correspond to sizes of the spore-derived colonies grown 3 days at 32° according to the spore genotype and the absence/presence of CFVII. For each medium and temperature, sizes were normalized by setting the mean size of the WT without CFVII at 100.

**S11 Fig.**
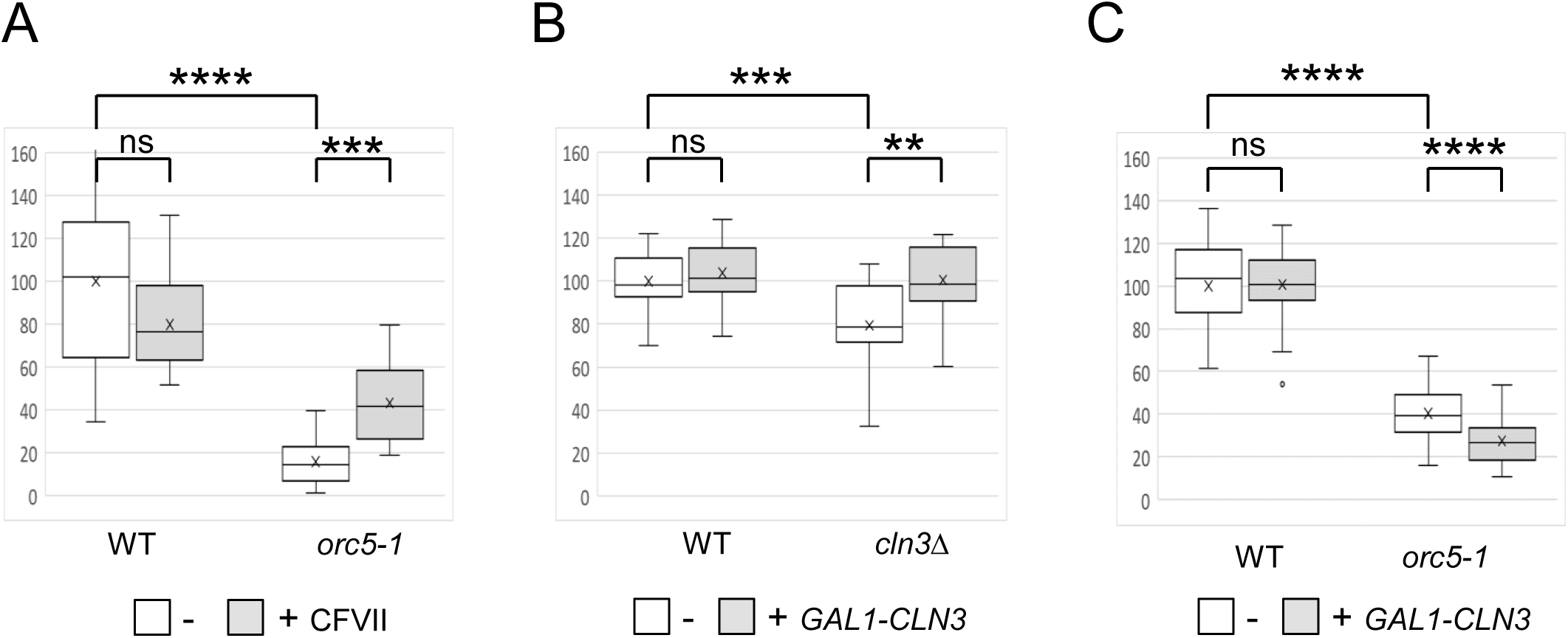
Effect of *CLN3* overexpression on *orc5-1* spore colony size. (A) Analysis of the meiotic products of the *ORC5/orc5-1* diploid with CFVII. The box plots correspond to sizes of the spore-derived colonies grown 4 days at 33,5° on YPGal according to the spore genotype and the presence of CFVII. Sizes were normalized by setting the mean size of the WT without CFVII at 100. P-values in a one-way ANOVA test are indicated : ns *p* > 0.05, * *p* < 0.05, ** *p* < 0.01, *** *p* < 0.001, **** *p* < 0.0001. (B) Same analysis as in (A) with the meiotic products of the *ORC5/orc5-1 GAL1-CLN3* diploid. The box plots correspond to sizes of the spore-derived colonies grown 3 days at 33,5° according to the spore genotype and the absence/presence of *GAL1-CLN3*. Sizes were normalized by setting the mean size of the WT without *GAL1-CLN3* at 100. (C) Same analysis as in (A) with the meiotic products of *ORC5/orc5-1 GAL1-CLN3* diploids. The box plots correspond to sizes of the spore-derived colonies grown 3 days at 34,5° on YPGal according to the spore genotype and the absence/presence of *GAL1-CLN3*. Sizes were normalized by setting the mean size of the WT without *GAL1-CLN3* at 100.

**S12 Fig.**
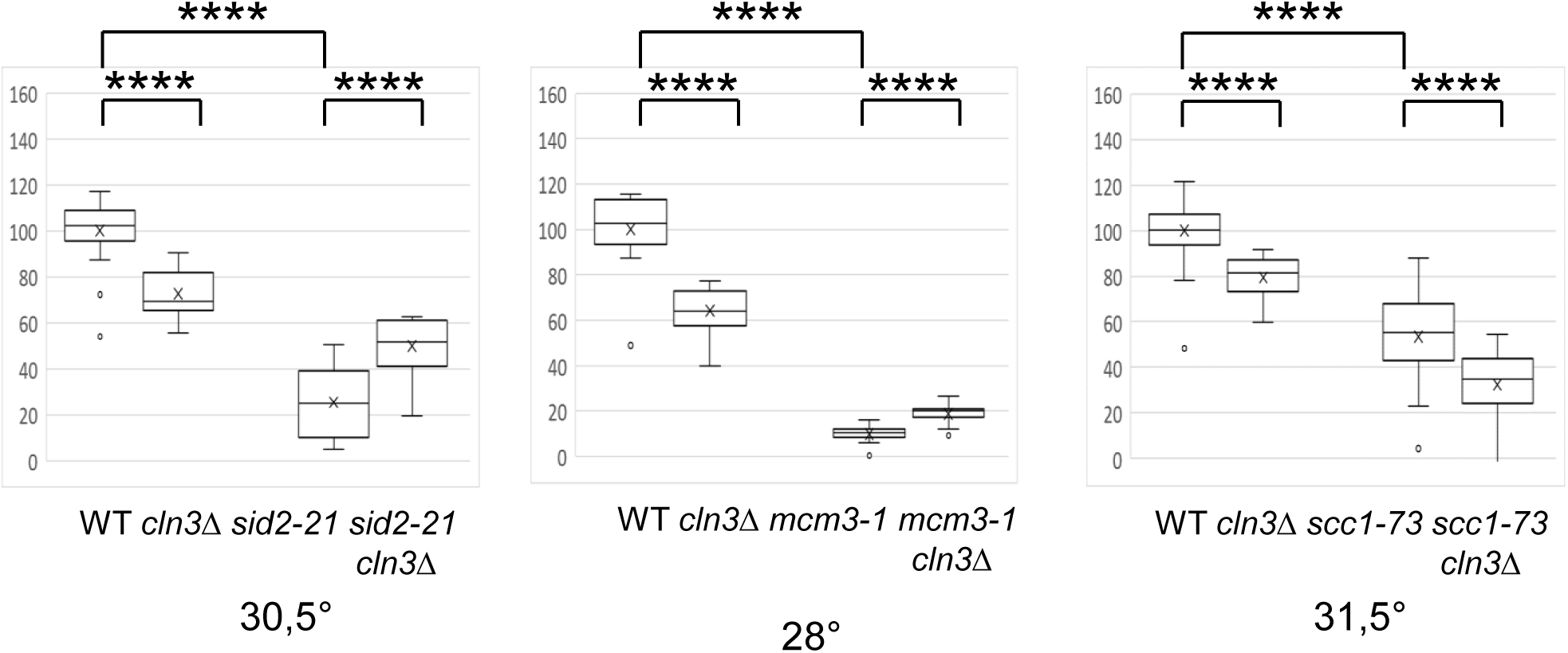
Effect of *cln3Δ* on *sid2-21*, *mcm3-1* and *scc1-73* spore colony size. Analysis of the meiotic products of the *SID2/sid2-21 CNL3/cln3Δ* (left), *MCM3/mcm3-1 CNL3/cln3Δ* (middle) and *SCC1/scc1-73 CNL3/cln3Δ* (right) diploids. The box plots correspond to sizes of the spore-derived colonies grown 2-3 days at the indicated temperature according to the spore genotype. Sizes were normalized by setting the mean size of the WT at 100. P-values in a one-way ANOVA test are indicated : ns *p* > 0.05, * *p* < 0.05, ** *p* < 0.01, *** *p* < 0.001, **** *p* < 0.0001.

**S13 Fig.**
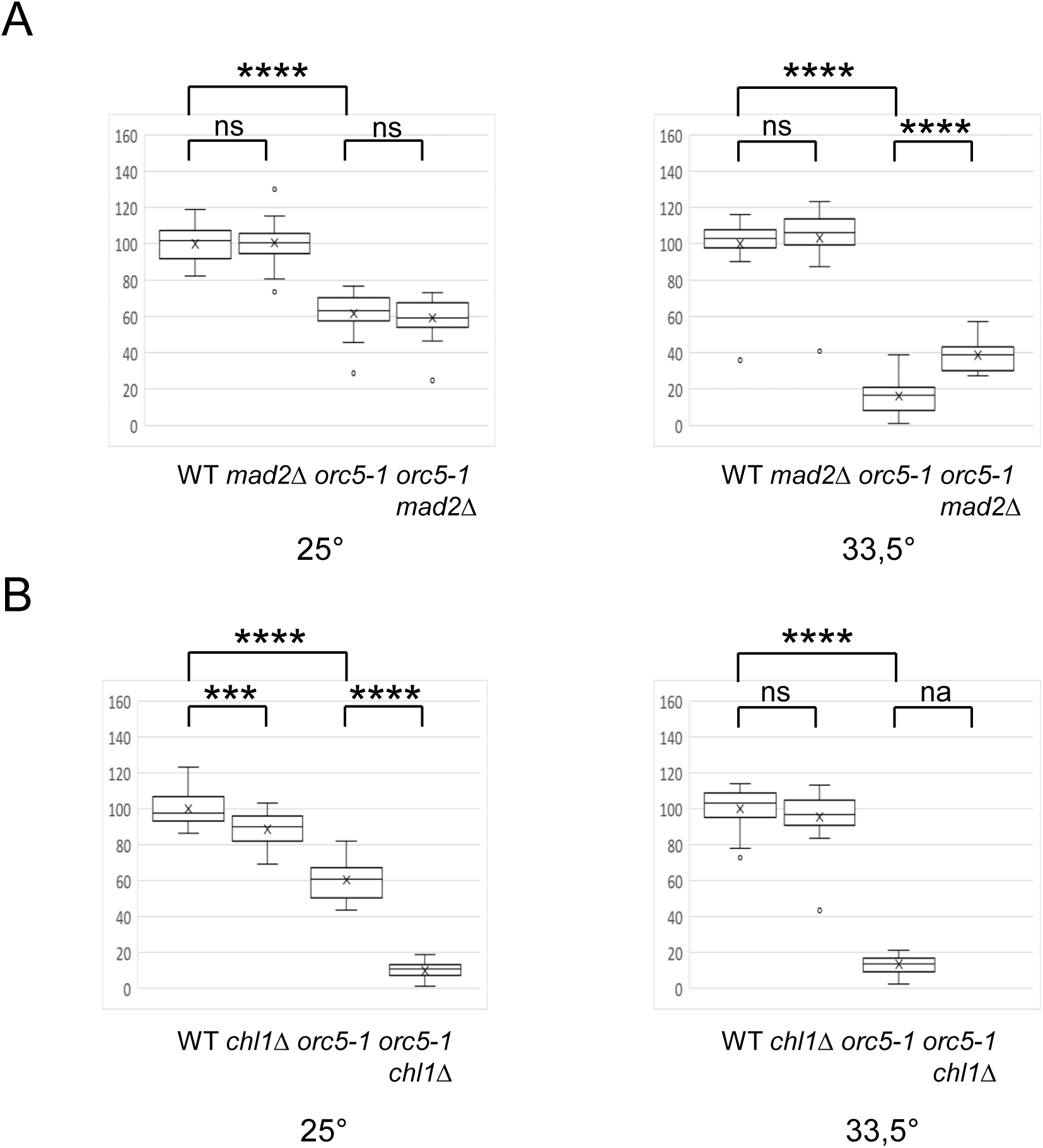
Opposite effects of *mad2Δ* and *chl1Δ* on *orc5-1* spore colony size. (A) Analysis of the meiotic products of the *ORC5/orc5-1 MAD2/mad2Δ* diploid. The box plots correspond to sizes of the spore-derived colonies grown 3 days at the indicated temperature according to the spore genotype. Sizes were normalized by setting the mean size of the WT at 100. P-values in a one-way ANOVA test are indicated : ns *p* > 0.05, * *p* < 0.05, ** *p* < 0.01, *** *p* < 0.001, **** *p* < 0.0001. (B) Same analysis as in (A) with the meiotic products of the *ORC5/orc5-1 CHL1/chl1Δ* diploid. na : not applicable.

**S14 Fig.**
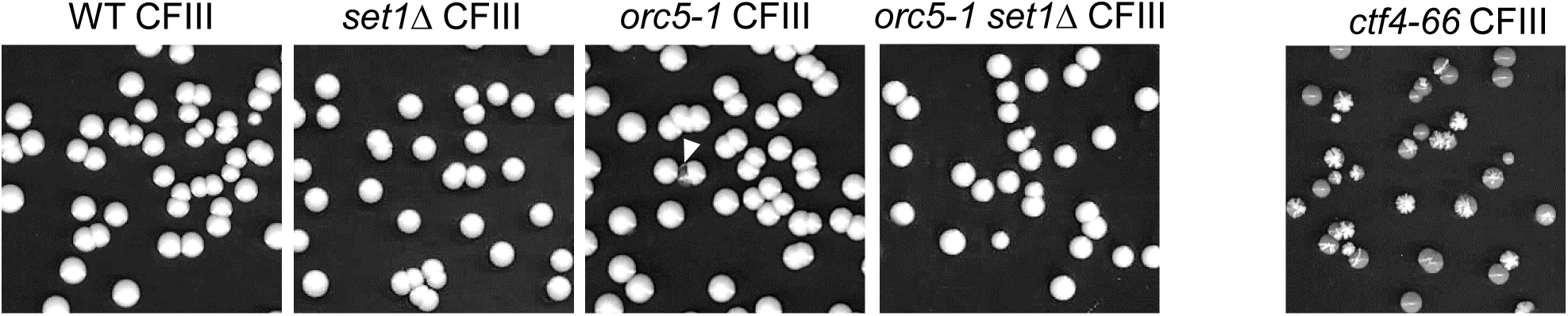
The *orc5-1* mutation is not associated with any detectable chromosomal instability. The images show yeast colonies of the wild type (*WT*) and mutant strains (as indicated) with CFIII after growth on YPD medium at 30°. The *ade2-1* mutation is responsible for the red color. The white color is due to the presence of CFIII. WT and mutant haploids were meiotic products of the same *ORC5/orc5-1 SET1/set1Δ* diploid with CFIII. A *ctf4-66* strain was used as a positive control. The white arrowhead indicates a red colony in *orc5-1* CFIII.

| Strains cerevisiae | Mating type | Genotype | Source |
| --- | --- | --- | --- |
| <u>SK1 background</u> |  |  | a - Valérie Borde<br>b - laboratory stock<br>c - Michael Weinreich |
| ORD5740 | a/α | *ura3/ura3 lys2/lys2 ho::LYS2/ho::LYS2 leu2/LEU2 his4/HIS4 arg4-nsp/arg4-bgl trp1/trp1* | a |
| CDK1120 | a/α | ORD5740 *orc5-1/orc5-1 set1::KanMX/set1::KanMX* | b |
| CDK1121 | a/α | ORD5740 *orc5-1/orc5-1 set1::KanMX/set1::KanMX rer1::HphMX/rer1::NatMX* | b |
| CDKA1 | a | *ura3 lys2 ho::LYS2 leu2 HIS4 arg4 trp1* | b |
| CDKA3 | α | *ura3 lys2 ho::LYS2 LEU2 his4 arg4 trp1* | b |
| CDKA11 | a | CDKA1 *rer1::NatMX* | b |
| CDKA12 | a | CDKA1 *rer1::HphMX* | b |
| CDKA111 | a | CDKA1 *rer1::NatMX/rer1::NatMX* (disomic for III) | b |
| CDKA312 | α | CDKA3 *rer1::HphMX/rer1::NatMX* (disomic for III) | b |
| <u>W303 background</u> |  |  |  |
| W303-1A | a | *ura3-1 leu2-3,-112 his3-11,-15 trp1-1 ade2-1 can1-100* | c |
| M197 | a | *ura3-1 leu2-3,-112 his3-11,-15 trp1-1 ade2-1 can1-100 orc5-1* | c |
| CDW1 | a | *ura3-1 leu2-3,-112 his3-11,-15 trp1-1 ade2-1 can1-100 set1::KanMX* | b |
| CDW2 | a/α | *ura3-1/ura3-1 leu2-3,-112/leu2-3,-112 his3-11,-15/his3-11,-15 trp1-1/trp1-1 ade2-1/ade2-1 can1-100/can1-100* | b |
| CDW20 | a/α | CDW2 *rer1::HphMX/rer1::NatMX/rer1::NatMX* (trisomic for III) | b |
| CDW201 | a/α | CDW20 *ORC5/orc5-1 SET1/set1::KanMX MRC1/MRC1/mrc1::TRP1* (trisomic for III) | b |
| CDW202 | a/α | CDW20 *ORC5/orc5-1 SET1/set1::KanMX DCC1/DCC1/dcc1::TRP1* (trisomic for III) | b |
| CDW2113 | a/α | CDW2 *ORC5/orc5-1 SET1/set1::KanMX CFIII-URA3-SUP11* | b |
| CDW2117 | a/α | CDW2 *ORC5/orc5-1 SET1/set1::KanMX CFVII-URA3-SUP11* | b |
| CDW21112 | a/α | CDW2 *ORC5/orc5-1 SET1/set1::KanMX ADE16/ade16::HIS/ade16::Kan* (trisomic for XII) | b |
| CDW223 | a/α | CDW2 *ORC5/orc5-1 RAD52/rad52::TRP1 CFVII-URA3-SUP11* | b |
| CDW217 | a/α | CDW2 *ORC5/orc5-1 CFVII-URA3-SUP11* | b |
| CDW21Y2 | a/α | CDW2 *ORC5/orc5-1 YAC-2 (yOX39)* | b |
| CDW21Y3 | a/α | CDW2 *ORC5/orc5-1 YAC-3 (yOX32)* | b |
| CDW37 | a/α | CDW2 *CFIII-URA3-SUP11 CFVII-URA3-SUP11* | b |
| CDW2137 | a/α | CDW2 *ORC5/orc5-1 CFIII-URA3-SUP11 CFVII-URA3-SUP11* | b |
| CDW227 | a/α | CDW2 *ORC2/orc2-1 CFVII-URA3-SUP11* | b |
| CDW267 | a/α | CDW2 *CDC6/cdc6-4 CFVII-URA3-SUP11* | b |
| CDW277 | a/α | CDW2 *CDC7/cdc7-1 CFVII-URA3-SUP11* | b |
| CDW2177 | a/α | CDW2 *CDC17/cdc17-1 CFVII-URA3-SUP11* | b |
| CDW2597 | a/α | CDW2 *SMC1/smc1-259 CFVII-URA3-SUP11* | b |
| CDW2427 | a/α | CDW2 *SMC3/smc3-42 CFVII-URA3-SUP11* | b |
| CDW2247 | a/α | CDW2 *SCC2/scc2-4 CFVII-URA3-SUP11* | b |
| CDW2147 | a/α | CDW2 *CDC14/cdc14-3 CFVII-URA3-SUP11* | b |
| CDW2157 | a/α | CDW2 *CDC15/cdc15-2 CFVII-URA3-SUP11* | b |
| CDW237 | a/α | CDW2 *SIC1/sic1::HIS3 CFVII-URA3-SUP11* | b |
| CDW213 | a/α | CDW2 *ORC5/orc5-1 CLN3/cln3::HIS3* | b |
| CDW2143 | a/α | CDW2 *CDC14/cdc14-3 CLN3/cln3::HIS3* | b |
| CDW273 | a/α | CDW2 *CDC7/cdc7-1 CLN3/cln3::HIS3* | b |
| CDW2173 | a/α | CDW2 *CDC17/cdc17-1 CLN3/cln3::HIS3* | b |
| CDW2103 | a/$\alpha$ | CDW2 *ORC5/orc5-1 GAL1-CLN3* | b |
| CDW2303 | a/$\alpha$ | CDW2 *CLN3/cln3::HIS3 GAL1-CLN3* | b |
| CDW2213 | a/$\alpha$ | CDW2 *SID2/sid2-21 CLN3/cln3::HIS3* | b |
| CDW233 | a/$\alpha$ | CDW2 *MCM3/mcm3-1 CLN3/cln3::HIS3* | b |
| CDW2733 | a/$\alpha$ | CDW2 *SCC1/scc1-73 CLN3/cln3::HIS3* | b |
| CDW2137 | a/$\alpha$ | CDW2 *ORC5/orc5-1 CLN3/cln3::HIS3 CFVII-URA3-SUP11* | b |
| CDW212 | a/$\alpha$ | CDW2 *ORC5/orc5-1 MAD2/mad2::TRP1* | b |
| CDW211 | a/$\alpha$ | CDW2 *ORC5/orc5-1 CHL1/chl1::HIS3* | b |
| CDW2127 | a/$\alpha$ | CDW2 *ORC5/orc5-1 MAD2/mad2::TRP1 CFVII-URA3-SUP11* | b |
| CDW2123 | a/$\alpha$ | CDW2 *ORC5/orc5-1 MAD2/mad2::TRP1 CLN3/cln3::HIS3* | b |

## Notes

### Competing Interest Statement

The authors have declared no competing interest.

